# Prophage-encoded phage defense proteins with cognate self-immunity

**DOI:** 10.1101/2020.07.13.199331

**Authors:** Siân V. Owen, Nicolas Wenner, Charles L. Dulberger, Ella V. Rodwell, Arthur Bowers-Barnard, Natalia Quinones-Olvera, Daniel J. Rigden, Eric J. Rubin, Ethan C. Garner, Michael Baym, Jay C. D. Hinton

## Abstract

Temperate phages are pervasive in bacterial genomes, existing as vertically-inherited islands called prophages. Prophages are vulnerable to the predation of their host bacterium by exogenous phages. Here we identify BstA, a novel family of prophage-encoded phage defense proteins found in diverse Gram-negative bacteria. BstA drives potent suppression of phage epidemics through abortive infection. During lytic replication, the *bstA*-encoding prophage is not itself inhibited by BstA due to a self-immunity mechanism conferred by the anti-BstA (*aba*) element, a short stretch of DNA within the *bstA* locus. Inhibition of phage replication by distinct BstA proteins from *Salmonella, Klebsiella* and *Escherichia* prophages is functionally interchangeable, but each possesses a cognate *aba* element. The specificity of the *aba* element ensures that immunity is exclusive to the replicating prophage, and cannot be exploited by heterologous BstA-encoding phages. BstA allows prophages to defend host cells against exogenous phage attack, without sacrificing their own lytic autonomy.

## Introduction

The perpetual battle between bacteria and their viruses (phages) has driven the evolution of a diverse array of phage defense systems in bacteria (Bernheim and Sorek, 2020; Hampton et al., 2020; Houte et al., 2016; Rostøl and Marraffini, 2019. Conversely, it is increasingly recognised that phages have evolved many mechanisms to subvert these defense systems (Maxwell, 2017; Samson et al., 2013; Trasanidou et al., 2019). Although the most intuitive form of phage defense might involve the direct rescue of an infected cell, for example by the targeted degradation of phage nucleic acids by CRISPR-Cas, or restriction modification systems, many phage-defense systems in fact function solely at the population level.

In a mechanism conceptually analogous to the pathogen-stimulated programmed cell death driven by the innate immune systems of higher organisms (Abedon, 2012), phage infection can be prevented from sweeping across populations, at the cost of the lives of infected cells. These population-level phage defense systems are often grouped under the umbrella term “abortive infection” (Abi) (Labrie et al., 2010; Lopatina et al., 2020) but actually represent diverse mechanisms to prevent phage replication and induce cell death. These include protease-mediated inhibition of cellular translation (Bingham et al., 2000), toxin-antitoxin pairs (Fineran et al., 2009; Pecota and Wood, 1996) and cyclic oligonucleotide signalling (Cohen et al., 2019). Recently, it has been proposed that certain CRISPR-Cas systems function through abortive infection (Meeske et al., 2019; Watson et al., 2019). Such mechanistic diversity and prevalence of abortive infection systems in bacteria emphasises the selective advantage this strategy imparts in the battle against phages.

However, an important sub-plot in the bacteria-phage conflict is the pervasive existence of so-called “temperate” or “lysogenic” phages *within* bacterial genomes. Temperate phages stably exist within the bacterial chromosome as latent, vertically-inherited islands known as a prophages. Crucially, to find new hosts, prophages must eventually escape from the bacterial genome and return to the lytic life-cycle. The prophage-state imposes unique existential pressures, wherein the fitness of the phage is indefinitely tied to that of the host bacterium. To indirectly enhance their own fitness, prophages frequently encode “moron” or “accessory” loci that modulate the biology of host bacteria (Bondy-Denomy and Davidson, 2014; Cumby et al., 2012; Fortier and Sekulovic, 2013; Howard-Varona et al., 2017), a phenomenon likened to altruism (Shub, 1994). Prophage accessory loci are often associated with bacterial virulence, and many notorious bacterial pathogens rely on prophage-encoded toxins and virulence factors to cause disease (Brüssow et al., 2004; Fortier and Sekulovic, 2013). However, another trait conferred by prophages that can significantly increase bacterial fitness is resistance against bacteriophage attack. Indeed, recent work has suggested that prophage accessory genes may represent an underexplored reservoir of phage-defense systems (Bondy-Denomy and Davidson, 2014; Dedrick et al., 2017; Snyder, 1995).

Here, we report a novel phage defense system driven by the BstA protein that is itself encoded by prophages of diverse Gram-negative bacteria. When a bacterium harbours a *bstA-*encoding prophage, the BstA protein confers effective population-level defense against exogenous phage via abortive infection. The *bstA* locus includes an anti-BstA element, which can suppress the activity of BstA protein to allow the native prophage to switch to a lytic lifestyle. We propose that this defense mechanism has evolved to allow prophages to both defend host cells from predatory phages and permit their own lytic replication.

## Results

### The BTP1 prophage-encoded *bstA* gene mediates phage resistance

*Salmonella enterica* subsp. *enterica* serovar Typhimurium (hereafter *S.* Typhimurium) strain D23580 encodes the ~40 kb prophage BTP1 (Figure 1A) (Owen et al., 2017). An operon encoded by BTP1, the *gtr* locus (*gtrAC^BTP1^*), confers resistance against phage P22 by chemically modifying the cellular lipopolysaccharide (LPS), the receptor for phage P22 (Kintz et al., 2015). Unsurprisingly therefore, deleting the BTP1 prophage from strain D23580 (D23580 ΔBTP1) made the strain highly susceptible to infection by phage P22, confirming that resistance to phage P22 is conferred by the BTP1 prophage (Figure 1B). However, inactivation of the *gtr* locus of prophage BTP1 (D23580 *Δtsp-gtrAC^BTP1^*) did not restore sensitivity to phage P22 to the level of D23580 ΔBTP1 (Figure 1B), suggesting the existence of a second BTP1-encoded phage resistance system.

**Figure 1:**
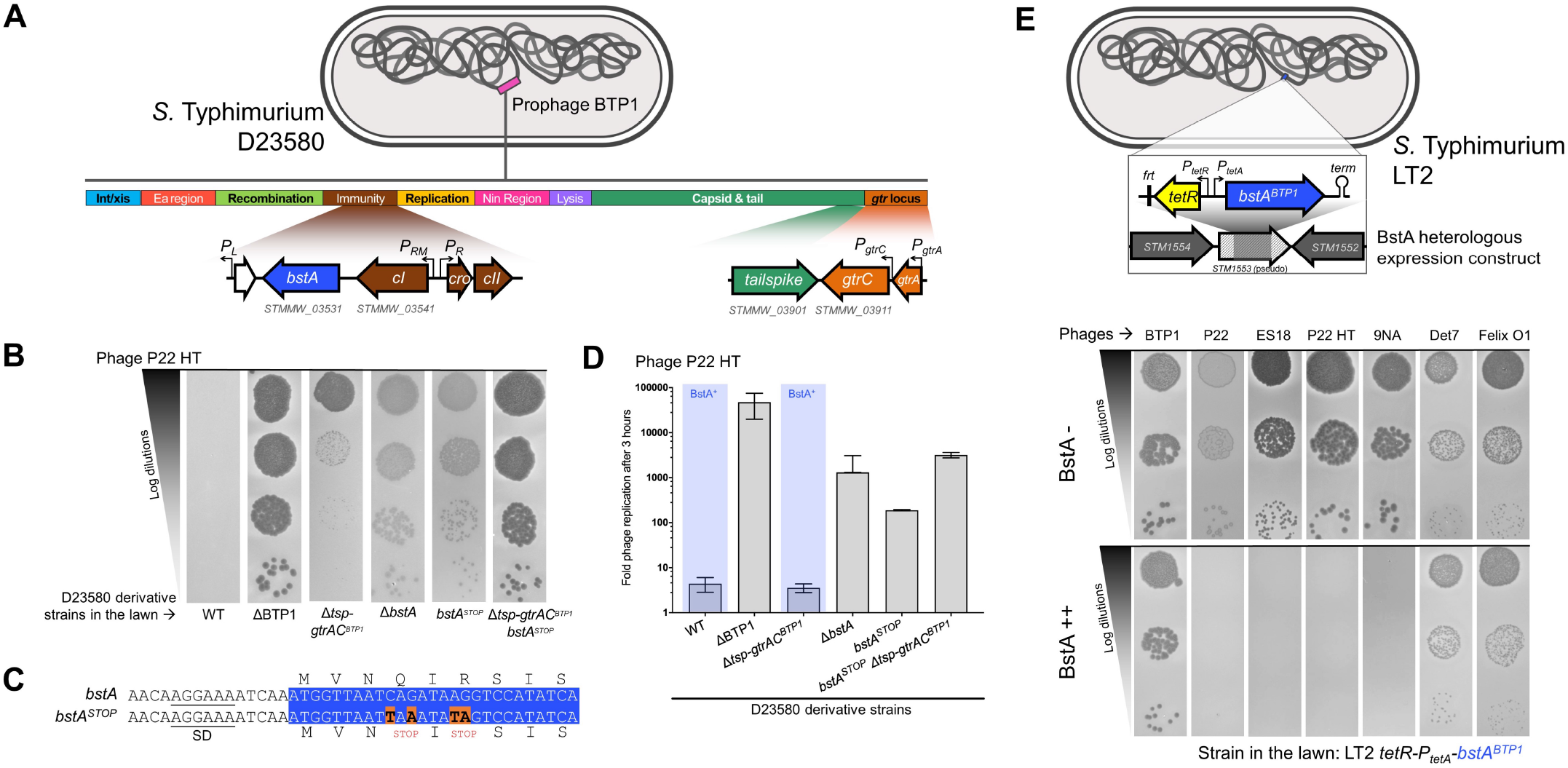
The *bstA* gene of prophage BTP1 confers phage defense. (**A**) Genomic architecture of prophage BTP1 of *S.* Typhimurium D23580, according to Owen *et al.*, 2020: the LPS modification genes *gtrAC^BTP1^* (Kintz et al., 2015) and the immunity region carrying *bstA* (downstream of the *cI* repressor gene) are detailed. Bent arrows represent promoters. For reference purposes the locus tags of important genes in this study in the D23580 reference genome (accession: FN424405) are shown. (**B**) Removal of prophage BTP1 from strain D23580 results in enhanced sensitivity to phage P22. Two BTP1 genes confer resistance to P22; *gtrAC* and *bstA*. Plaque assays were performed with phage P22 HT *105/1 int-201* (P22 HT) applied to lawns of *S.* Typhimurium D23580 WT or ΔBTP1, *ΔbstA, bstA^STOP^, Δtsp-gtrAC^BTP1^* and *Δtsp-gtrAC^BTP1^ bstA^STOP^* mutants (strains JH3877, SSO-204, SSO-78, JH4287 and SNW431, respectively). The requirement for the inactivation of *tsp* is describes in the Methods. (**C**) The 4 nucleotide substitutions leading to two nonsense mutations in the *bstA^STOP^* strain are indicated. SD: putative Shine-Dalgarno sequence of the *bstA* gene. The beginning of the *bstA* open reading frame is highlighted in blue. (**D**) Phage replication assays in liquid culture using P22 HT and the same D23580 derivative strains shown in the plaque assay in B. Replication was measured 3 hours post-infection and phages were enumerated on lawns of D23580 *Δtsp-gtrAC bstA^STOP^* (SNW431). Phage replication is presented as the mean of biological triplicates ± SD. (**E**) BstA protein confers phage defense in *S.* Typhimurium LT2. Phages P22, ES18, P22HT and 9NA are inhibited by BstA. Phages Det7, Felix O1 and BTP1 are not affected by BstA expression. Plaque assays were carried out with the indicated *Salmonella* phages applied to lawns of LT2 *tetR-P_tetA_-bstA^BTP1^* (JH4400) in the absence (BstA-) or the presence of the inducer anhydrotetracycline (AHT, BstA++). The *tetR-P_tetA_-bstA* insertion replacing a part of the *STM1553* pseudogene of strain JH4400 is schematized above: *tetR* encodes the tetracycline repressor that represses the *P_tetA_* promoter in the absence of AHT induction, *“frt”* denotes the 84 nt scar sequence of pKD4 and the hairpin represents the native *bstA* Rho-independent terminator (*term*).

Our previous transcriptomic study showed that the *bstA* gene was highly-expressed from prophage BTP1 during lysogeny, making it a candidate novel phage accessory gene (Owen et al., 2020). The *bstA* gene, encoded downstream of the prophage *cI* repressor locus, has been implicated phenotypically in both virulence and anti-virulence of *Salmonella* isolates, but no functional mechanism has been proposed (Herrero-Fresno et al., 2014, 2018; Spiegelhauer et al., 2020), and the BstA protein has not been characterized. We hypothesised that *bstA* was the second element in the BTP1 prophage that conferred defense against phage P22.

Consistent with this hypothesis, removal of the *bstA* gene from prophage BTP1 (D23580 *ΔbstA*) dramatically increased susceptibility to phage P22 (Figure 1B). To confirm that phage resistance was directly mediated by BstA protein, we introduced two stop codons into the beginning of the *bstA* coding sequence by exchanging 4 nucleotides (D23580 *bstA^STOP^*) (Figure 1C). D23580 *bstA^STOP^* was highly susceptible to P22 phage, to the same level as D23580 *ΔbstA*, demonstrating that BstA protein mediates defense against phage P22. Simultaneous deletion of the *gtr* locus and inactivation of the BstA protein (D23580 *Δtsp-gtrAC^BTP1^ bstA^STOP^*) recapitulated the susceptibility to phage P22 achieved by deleting the entire BTP1 prophage (D23580 ΔBTP1), indicating that resistance to phage P22 was solely mediated by the *bstA* and *gtrAC* loci in prophage BTP1. The findings were reproduced by assaying the replication of phage P22 on the same strains in liquid culture, demonstrating quantitatively that reduction of plaque formation by BstA truly reflected suppression of phage replication (Figure 1D).

To investigate whether the defense function of the BstA protein depended on any other elements from the BTP1 prophage, we constructed an inducible expression system in *S.* Typhimurium strain LT2 which does not contain the BTP1 prophage. LT2 is the type strain of *S.* Typhimurium, and is natively susceptible to many phages, including P22 (McClelland et al., 2001). Expression of the BstA^BTP1^ protein in *S.* Typhimurium LT2 from within a neutral position on the chromosome (LT2 *tetR-P_tetA_-bstA*) conferred a high degree of resistance to P22 and other phages, including ES18 and 9NA (Figure 1E; Supplementary Figure 1A).

Whilst induced expression of BstA^BTP1^ completely eliminated plaque formation of sensitive phages, at very high phage concentrations (10^9-10^ PFU/mL) these phages still produced some clearing of the bacterial lawn (Supplementary Figure 1B), consistent with some mechanisms of phage defense, such as abortive infection. Expression of the derivative containing two stop codons at the beginning of the *bstA* coding sequence (*bstA^STOP^*) conferred no phage resistance, demonstrating again that defense is mediated by *bstA* at the protein level (Supplementary Figure 1C). However, BstA did not mediate resistance against all phages tested: Det7, Felix O1, and notably, phage BTP1 (which encodes the *bstA* gene) were unaffected by expression of BstA, both at the level of plaque assay and replication in liquid culture (Figure 1E; Supplementary Figure 1A). Induction of *bstA* or *bstA^STOP^* expression in the absence of phage infection did not cause any detectable effect on cell growth rate, suggesting that overexpression of BstA^BTP1^ does not cause toxicity (Supplementary Figure 1D). We were unable to detect any pattern in the characteristics or gene repertoire of phages that were sensitive or insensitive to BstA protein that could relate to the mechanistic action of BstA protein.

### BstA represents a novel family of prophage-encoded phage defense proteins in diverse Gram-negative bacteria

Having established that BstA functions as a prophage-encoded phage defense system, we sought to further characterise the evolutionary conservation of this protein. We identified BstA homologs in the genomes of diverse Gram-negative bacteria (Supplementary Table 1) and compiled a dataset of 72 homologs representative of phylogenetic diversity. The majority (79%) of these BstA homologs co-occurred with phage genes, and were designated as putatively-prophage associated (Figure 2A). No known phage-associated genes were found in the vicinity of 21% (15 of 72) of BstA homologs, which were defined as putatively prophage-independent. A small subset of BstA homologs were plasmid-encoded, a group that included both putatively prophage-associated and prophage-independent homologs (Figure 2A). Strikingly, in many cases, BstA homologs were located downstream of putative prophage repressor proteins, mirroring the genetic architecture of BstA^BTP1^ (Figure 2B). We conclude that the BstA protein is highly associated with prophages of Gram-negative bacteria.

**Figure 2:**
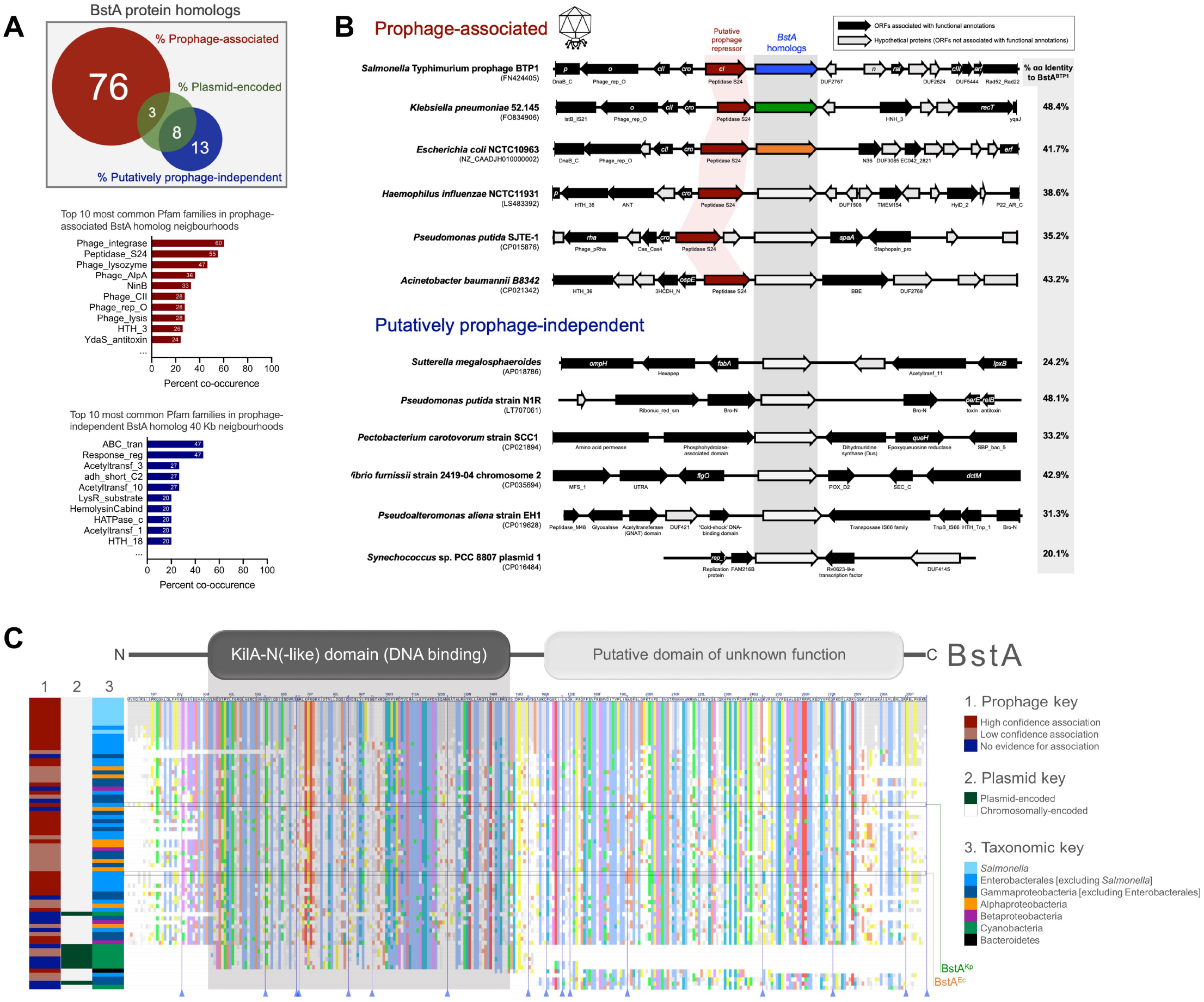
BstA homologs are found in diverse bacterial taxa and are frequently associated with prophages. (**A**) A dataset of 72 BstA homologs representative of taxonomic diversity were manually curated and analyzed for prophage association based on the co-occurrence of phage-related Pfam-domains in the 20 kb either side of each homologs (yielding a total 40 kb window) (Supplementary Table 1). Homologs without co-occurring phage-related protein domains were assigned to be “Putatively prophage-independent”. A further subset of the BstA homologs were encoded on plasmids. The top ten most commonly co-occurring Pfam domains with prophage-associated and putatively prophage-independent BstA homologs are shown as bar graphs. (**B**) Gene maps showing the genetic context of a selection of 6 prophage-associated and 6 putatively prophage-independent BstA homologs (homologs indicated by the grey rectangle). Putative prophage repressor genes are highlighted in red. The top three BstA proteins from BTP1 (BstA^BTP1^, blue), *K. pneumoniae* 52.145 (BstA^Kp^, green) and *E. coli* NCTC10963 (BstA^Ec^, orange) are studied experimentally in later stages of this work, and so are highlighted. Open reading frames associated with functional annotations are shown as solid black arrows, and functional gene name or Pfam domains are annotated. (**C**) An alignment of the 72 BstA protein homologs to BstA^BTP1^, with colors indicating amino acid conservation (Clustal colour scheme). Alignment columns containing gaps relative to the reference sequence (BstA^BTP1^) have been collapsed and are indicated with blue lines and triangles at the base of the alignment (an expanded alignment can be found in Supplementary Figure 2). The position of BstA^Kp^ and BstA^Ec^ within the alignment is highlighted. The position of the KilA-N (-like) domain (BstA^BTP1^ residues 32-147) is indicated by a grey box. Heatmaps on the left of the alignment indicate the prophage and plasmid association of each homolog (lanes 1 & 2), and the taxonomic group each homolog derives from (lane 3). Prophage association was split into high and low confidence based on gene co-occurrence criteria (see Methods).

Whilst the BstA protein does not exhibit sequence homology to any functionally-characterised proteins, remote homology detection methods revealed a KilA-N (-like) domain in the N-terminal region (residues 32-147 of BstA^BTP1^) (Figure 2C). Though poorly characterised, the KilA-N domain is found in proteins from phages and eukaryotic DNA viruses, and contains the helix-turn-helix motif characteristic of DNA binding proteins (Iyer et al., 2002; Medina et al., 2019). A large screen based on genomic proximity to known-phage defense systems previously implicated KilA-N domain containing proteins in phage defense (Doron et al., 2018). Though the KilA-N domain derives its name from the *kilA* gene product of bacteriophage P1 (lethal when expressed in *E. coli*), *kilA* has no known function in phage infection biology (Hansen, 1989), and no bacterial KilA-N domain-containing proteins have yet been functionally characterised.

Certain residues in the BstA protein are highly conserved amongst homologs from diverse members of the Alpha-, Beta-, and Gamma-Proteobacteria (Figure 2C; Supplementary Figure 2A). A small number of BstA protein homologs (all found in Cyanobacterial plasmids) only exhibited homology to the N-terminal, KilA-N (-like) domain. A second small group of homologs derived from Proteobacteria, Cyanobacteria and a single Bacteroidetes isolate were only homologous to the C-terminal region of BstA (shown at the bottom on the alignment in Figure 2C). Such bipartite protein homology suggests that the BstA protein is composed of two functional domains. This conclusion is independently supported by evolutionary covariance analysis (Supplementary Figure 2B) where the clear depletion of predicted residue contacts between the ranges 1 to ~155 and ~156-307 of BstA^BTP1^ suggests that there is a domain boundary (Rigden, 2002) around position 155, with the two folded domains making few contacts. Additionally, we observed that the identity of BstA protein homologs did not obviously correlate with bacterial taxonomy (and inferred broad phylogeny). For example, homologs from enteric bacteria closely-related to *Salmonella* (members of the Gammaproteobacteria) sometimes shared less amino acid identity to BstA^BTP1^ than homologs from alpha- and betaproteobacteria, suggesting BstA proteins may be horizontally transferred, perhaps consistent with their association with prophages.

We selected two diverse BstA homologs from *Klebsiella pneumoniae* (48.4% amino acid identity to BstA^BTP1^) and *E. coli* (41.7% amino acid identity) to investigate the phage-resistance function of the larger BstA protein family (the native genetic context of these homologs is illustrated in Figure 2B, and their identity to BstA^BTP1^ is highlighted in the alignment in Figure 2C). We engineered inducible expression systems mirroring the expression construct previously validated for BstA^BTP1^ (Figure 3A,B; Figure 1E). Expression of BstA^Kp^ and BstA^Ec^ in *S.* Typhimurium LT2 conferred resistance to *Salmonella* phages at a similar level to BstA^BTP1^, despite these BstA homologs only sharing around 40% identity at the amino acid level (Figure 3A,B; Supplementary Figure 3A). Importantly, BstA^Kp^ and BstA^Ec^ prevented replication of phage BTP1 (which encodes *bstA^BTP1^),* unlike BstA^BTP1^.

**Figure 3:**
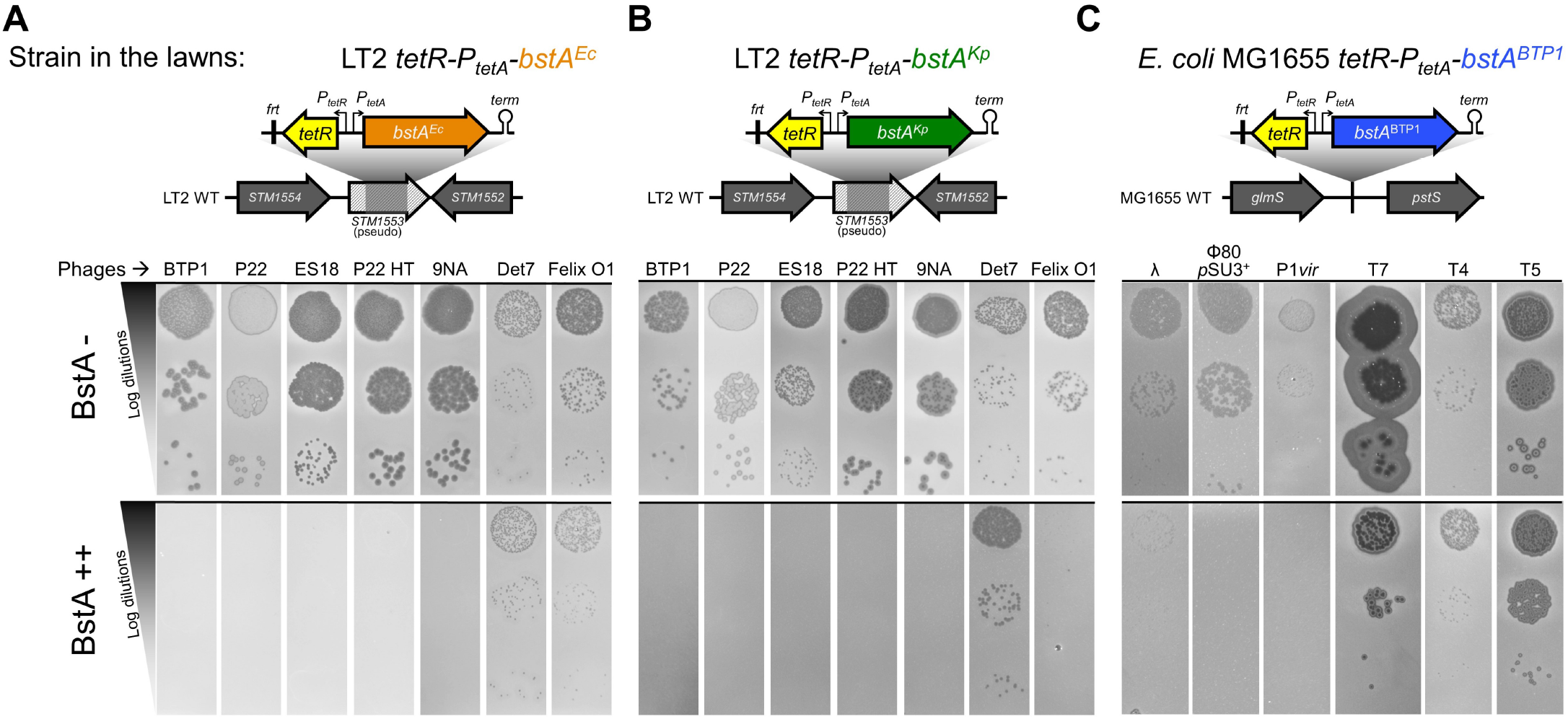
BstA homologs from *Salmonella, E. coli* and *K. pneumoniae* confer phage defense. Heterologous expression of *bstA* homologs from (**A**) *E. coli* NCTC10963 (*bstA^Ec^*) and (**B**) *K. pneumoniae* Kp52.145 (*bstA^Kp^*) in *Salmonella* strain LT2 confers phage defense at similar levels to *bstA^BTP1^*, but show additional activity against phage BTP1. (**C**) BstA^BTP1^ confers defense against coliphages in *E. coli* MG1655. Plaque assays were carried out with the indicated phages applied on mock-induced (BstA−) or AHT-induced (BstA++) lawns of LT2 *tetR-P_tetA_-bstA^Ec^* (JH4408), LT2 *tetR-P_tetA_-bstA^Kp^* (JH4404) or MG1655 *tetR-P_tetA_-bstA^BTP1^* (JH4410). The genetic context for the *tetR-P_tetA_-bstA* insertions within the *SM1553* pseudogene of LT2 or in the *glmS-pstS* intergenic region of MG1655 are depicted above.

Finally, to investigate the phage-defense function of BstA against well-characterised coliphages, we expressed BstA^BTP1^ and BstA^Ec^ in *E. coli*. Heterologous expression of BstA^BTP1^ in *E. coli* strain MG1655 conferred resistance to phage λ, Φ80, P1 and T7, but did not affect phages T4 and T5 (Supplementary Figure 3B,C). Surprisingly we found BstA^Ec^ was slightly less active against coliphages than BstA^BTP1^ (Supplementary Figure 3B,C). Replication in liquid culture was a more reliable and reproducible measure of phage susceptibility than plaque assay, and frequently revealed stronger resistance phenotypes than by plaque assay (Supplementary Figure 3B,C).

We conclude that BstA represents a novel family of phage-resistance proteins associated with the prophages of diverse Gram-negative bacteria.

### BstA mediates effective population-level phage defense through abortive infection

Phage resistance systems operate *via* diverse functional mechanisms (Hampton et al., 2020; Rostøl and Marraffini, 2019). We used a microscopy-based approach to dissect BstA-mediated phage-resistance. Virulent P22 phages (P22 Δ*c2*) were used to infect *Salmonella* cells with and without native BstA^BTP1^ function, at high multiplicity of infection (MOI) to ensure that most cells were infected. We were surprised to observe that independent of BstA^BTP1^ function, all cells lysed within the time course of 3 hours (Figure 4A, Supplementary Video 1), and BstA^BTP1^ function did not appear to confer any direct protection from phage infection at the level of individual infected cells. We conducted the same experiment in liquid culture, measuring phage replication and the fraction of surviving cells *post* phage infection. In cells possessing functional BstA (D23580 *Δtsp-gtrAC*), phage P22 *Δc2* completely failed to replicate (Figure 4B). In contrast, in the absence of BstA function (D23580 *Δtsp-gtrAC bstA^STOP^*), the phage replicated >100-fold. However, despite preventing the replication of phage P22, BstA^BTP1^ had no effect on cell survival: independent of BstA^BTP1^ function only 1-2% of cells survived following P22 infection (Figure 4C). We hypothesised that rather than rescuing infected cells, BstA protein must instead mediate phage defense at the population level.

**Figure 4:**
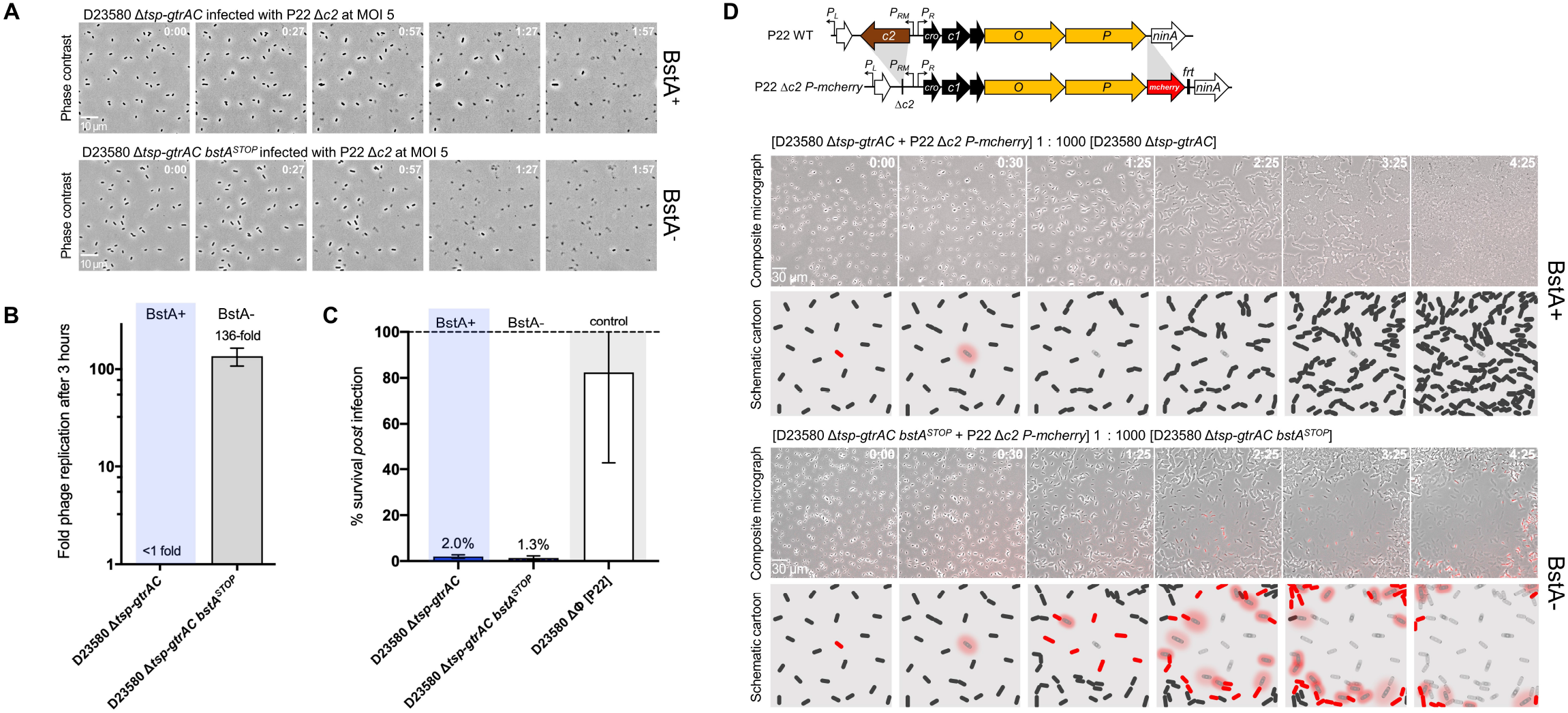
BstA mediates population-level phage defense through abortive infection. BstA protein does not protect individual cells from phage infection. (**A**) Cells natively expressing BstA (D23580 *Δtsp-gtrAC*, JH4287) or possessing a mutated BstA locus (D23580 *Δtsp-gtrAC bstA^STOP^*, SNW431) were infected with the obligately virulent P22-derivate phage, P22 *Δc2*, at an MOI of 5 to increase the likelihood of infecting all cells. Infected cells were imaged on agarose pads and the images represent a time series. Regardless of BstA function, almost all cells were observed to lyse (indicated by loss of defined cell shape and phase contrast). Videos of the time series are presented in Supplementary Video 1. (**B**) A phage replication assay showed that P22 Δ*c2* phage failed to replicate after three hours growth on the BstA+ strain (D23580 *Δtsp-gtrAC*), but replicated ~136-fold when BstA was inactivated (D23580 *Δtsp-gtrAC bstA^STOP^*). (**C**) Survival assay of the same strains after infection by phage P22 Δ*c2*, at an MOI of 5. Consistent with the microscopy data in (A), BstA function did not affect cell survival from phage infection. D23580 ΔΦ [P22] (SSO-128), a phage P22 lysogen (and therefore natively resistant), was used as a negative control. (**D**) A fluorescent reporter module for phage replication was added to P22 Δ*c2* (P22 Δ*c2 P-mcherry*) so that phage replication yielded red fluorescence (*mcherry* was inserted into the replicative genes of the phage). A similar experiment to (A) was conducted, but P22 Δ*c2 P-mcherry* infected cells were mixed 1:1000 with uninfected cells. In the BstA+ cells, primary infected cells lysed, but did not stimulate secondary infections of neighbouring cells, and eventually formed a confluent lawn. In BstA-cells, primary lysis events caused secondary infections (neighbouring cells showing red fluorescence and subsequent lysis) causing an epidemic of phage infection reminiscent of plaque formation. Cartoons schematise the outcomes of these experiments in the two strain backgrounds. Videos of the time series are presented in Supplementary Video 2. All experiments were carried out in liquid or solid M9 Glu^+^ media.

To investigate whether BstA protein mediated population-level phage defense, we conducted a second microscopy experiment, wherein approximately only 1 in every 1000 cells was infected with phage P22. Unlike culture in liquid media, our microscopy setup involved immobilisation of cells on agarose pads, which restricts the movement of phage particles to localized diffusion. The spread of infection was tracked as primary infected cells lysed and produced secondary infections. To visualise these phage epidemics, we used a reporter phage engineered to encode the red fluorescent protein mCherry within the early lytic operon (P22 *Δc2 P-mCherry*); the fluorescence signal indicated phage replication (Figure 4D).

In the population lacking functional BstA^BTP1^ (D23580 *Δtsp-gtrACbstA^STOP^*), primary infected cells lysed after around 30 minutes (Figure 4D, Supplementary Video 2). Subsequently, the red fluorescence signal was observed in neighbouring cells revealing secondary infection, followed by cell lysis, a cycle which repeated until all cells in the radius of the primary infected cell had lysed, reminiscent of plaque formation (Figure 4C). The impact of the epidemic of phage infection upon bacterial cells lacking BstA^BTP1^ can be visualised in Supplementary Video 2.

In contrast, native BstA activity prevented infection of the D23580 *Δtsp-gtrAC* population from generating a red fluorescence signal. Following lysis of the primary infected cells, no secondary infection was observed in neighbouring cells. Instead, cells continued to divide normally, eventually forming a confluent lawn (Figure 4D, Supplementary Video 2). The lack of subsequent rounds of secondary infection after the primary cell lysis events shows that few or no infectious phage particles were generated.

Taken together, these experiments demonstrate that BstA protein inhibits successful phage replication, but does not prevent the death of the infected cell. BstA therefore provides phage defense at the population-level and prevents the spread of phage epidemics. Accordingly, we propose that BstA is a novel abortive infection system: a population-level phage defense system that inhibits phage infection by sacrificing cell viability.

### BstA protein responds dynamically to phage infection and co-localises with phage DNA

To explore the molecular activity of BstA during phage infection, we first constructed a translational fusion of the BstA^BTP1^ protein to superfolder green fluorescent protein (_sf_GFP), and confirmed that the translational fusion did not compromise the function of the BstA protein (Supplementary Figure S4). We then used time-lapse fluorescence microscopy to observe the dynamics of BstA protein inside individual cells during infection with two BstA-sensitive phages, P22 and 9NA. In the absence of phage infection, BstA protein was distributed diffusely within the cytoplasm of the cells, suggesting no particular sub-cellular localisation (Figure 5A, Supplementary Video 3). However, approximately 20 minutes after infection with phages P22 and 9NA, we consistently observed BstA protein aggregating into discrete foci towards the centre of infected cells (Figure 5B, Supplementary Video 3). Cell lysis occurred approximately 40 minutes after the formation of BstA foci.

**Figure 5:**
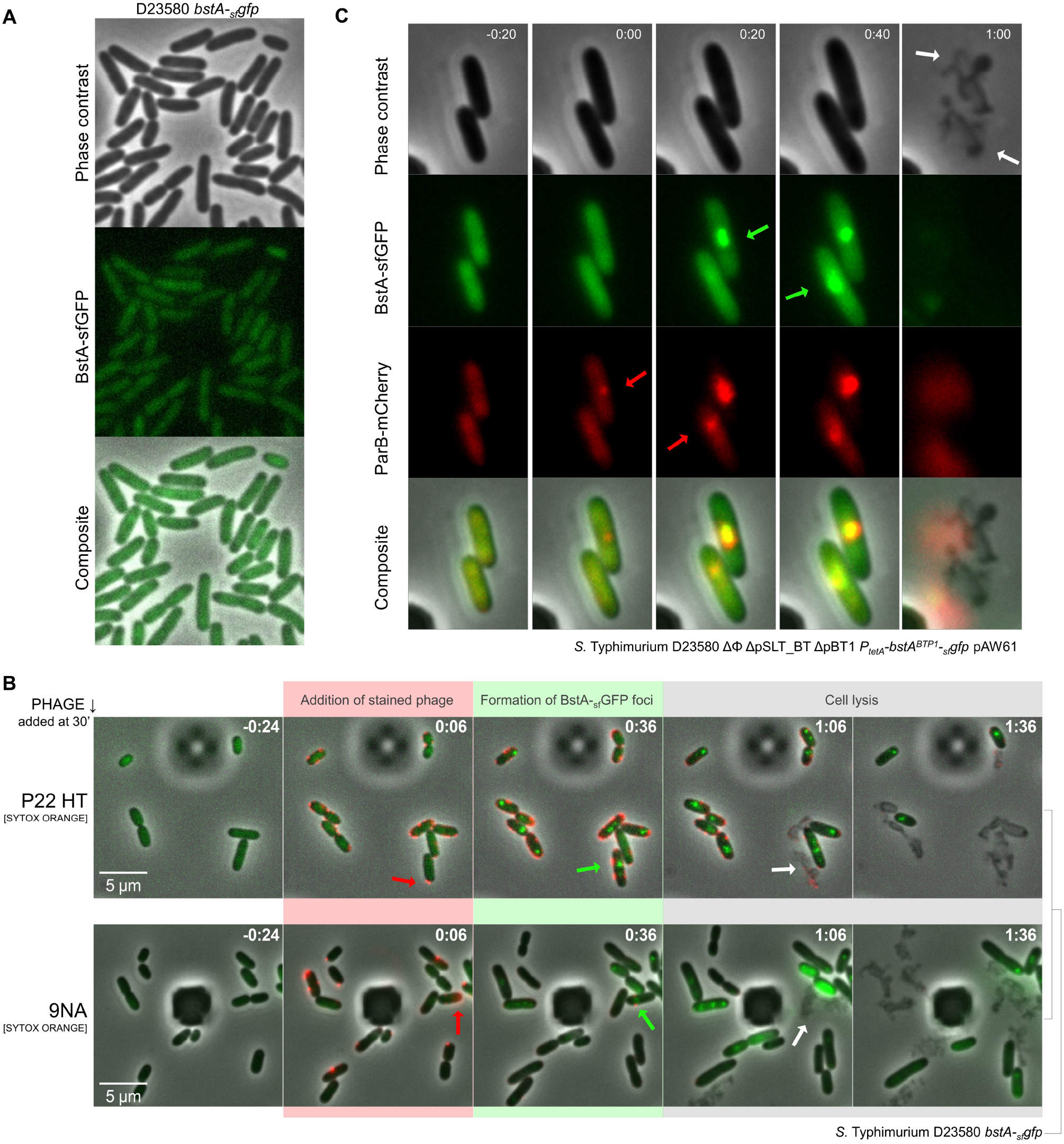
BstA protein responds dynamically to phage infection and colocalises with phage DNA. (**A**) A translational fusion of BstA to _sf_GFP protein (D23580 *bstA-_sf_gfp*, SNW403) was constructed to track the location of BstA protein inside cells. In the absence of phage infection, GFP signal is diffuse within the cell cytoplasm, suggesting no specific sub-cellular localisation. (**B**) A microfluidic growth chamber was used to observe the behaviour of BstA protein during phage infection, capturing images every 1.5 minutes. A time series of representative fields are presented as composite images (phase contrast, green and red fluorescence are overlaid). Cells were first grown for a period in the chamber (immobilised by the angle of the chamber ceiling) with constant flow of M9 Glu^+^ media (Methods). Fluorescently-labelled phages P22 HT or 9NA (stained with SYTOX Orange resuspended in M9 Glu^+^ media, Methods) were then added to the cells, and can be seen adsorbing to cells as red fluorescent puncta (red arrows). For comparative purposes, timestamps are synchronised to the point at which phage were first observed adsorbing to cells. Typically around 20 minutes after initial observation of phage infection, BstA proteins formed discrete and dynamic foci within the cells (green arrows). Cells then proceeded to lyse (white arrows), consistent with previous microscopy data in Figure 4. Videos of the time series are presented in Supplementary Video 3. (**C**) A microfluidic growth chamber was used to co-localise BstA protein and the DNA of the infecting phage. Prophage-free and plasmid-free SVO251 cells (expressing the BstA-_sf_GFP fusion and a ParB-mCherry fusion protein) were grown for a period of 15 minutes before P22 Δ*pid*::(*parS-aph*) phages were flowed across the cells. The ParB-mCherry fusion protein oligomerises at the *parS* site on the infecting phage chromosome, so that mCherry foci indicate the subcellular location of phage DNA (red arrows). To facilitate comparison, the time stamp was set to zero at the first observation of mCherry foci (i.e. the earliest detectable event of phage infection). BstA foci formed directly after appearance of mCherry foci (green arrows), and the merging of the images (composite) confirmed an overlap of the foci, consistent with the co-localisation of BstA foci with infecting phage DNA. Cells proceeded to lyse (white arrows). Time is indicated as h:m, and videos of the time series are presented in Supplementary Video 4.

We speculated that the dynamic establishment of foci by BstA in response to phage infection was likely to reflect the mechanistic activity of the protein. We noticed that the dynamics of the foci formed by BstA proteins during phage infection resembled live-cell fluorescence microscopy studies of phage replisomes (Cenens et al., 2013; Trinh et al., 2017). We therefore speculated that the focus of BstA protein in phage infected cells might correspond to the replicating phage DNA. To test this hypothesis, we used a ParB-*parS* system to track the sub-cellular localisation of phage DNA relative to BstA protein. We inserted a *parS* site into the P22 phage chromosome, and expressed a ParB-mCherry fusion protein inside cells already expressing BstA-_sf_GFP. ParB protein oligomerises onto DNA at *parS* sites, labelling *parS*-tagged DNA with ParB-mCherry foci. We conducted a microfluidic infection experiment to co-locate BstA foci and infecting P22 phage DNA, and observed that the position of ParB-mCherry foci (corresponding to phage P22 DNA) clearly overlapped with foci formed by BstA-_sf_GFP (Figure 5C, Supplementary Video 4). The microscopy data therefore suggest that BstA protein interacts with the replicating DNA of infecting phages. Consistent with the other microscopy data (Figure 4A, Figure 5B), cells proceeded to lysis after the formation of BstA/ParB-mCherry foci. We note that the strain used in this experiment (SVO251, Supplementary Table 2) is cured of all prophages, ruling out the possibility that cell lysis is caused by native prophage induction.

In summary, our data are consistent with a model that involves the movement of BstA protein to sites of phage DNA replication inside infected cells, followed by prevention of successful multiplication of the phage.

### BstA phage resistance systems contain anti-BstA elements (*aba*) that suppress the activity of BstA

When characterising the sensitivity of different phages to the activity of BstA^BTP1^ using our heterologous expression system (Figure 1E), we observed that BTP1 phage, (which itself encodes the *bstA^BTP1^* gene) was not affected by expression of BstA^BTP1^ (schematised in Figure 6A). We hypothesized that BTP1 carried an anti-BstA determinant: a self-immunity factor that allows phage BTP1 to replicate without being targeted by its own abortive infection protein. Consistent with this hypothesis, phage BTP1 became sensitive to BstA^BTP1^ expression when the *bstA* coding sequence was deleted (BTP1 *ΔbstA*). (Figure 6B). The self-immunity function of the *bstA* locus was not affected by the introduction of the double stop codon mutation into the beginning of the coding sequence (as described in Figure 1C), indicating that self-immunity is not mediated by the BstA protein itself, but by an alternative genetic element encoded within the *bstA* locus (Figure 6B). Here, and for the duration of this report, we define the *bstA* “locus” as the region including the *bstA* coding sequence with its 5’ upstream sequence.

**Figure 6:**
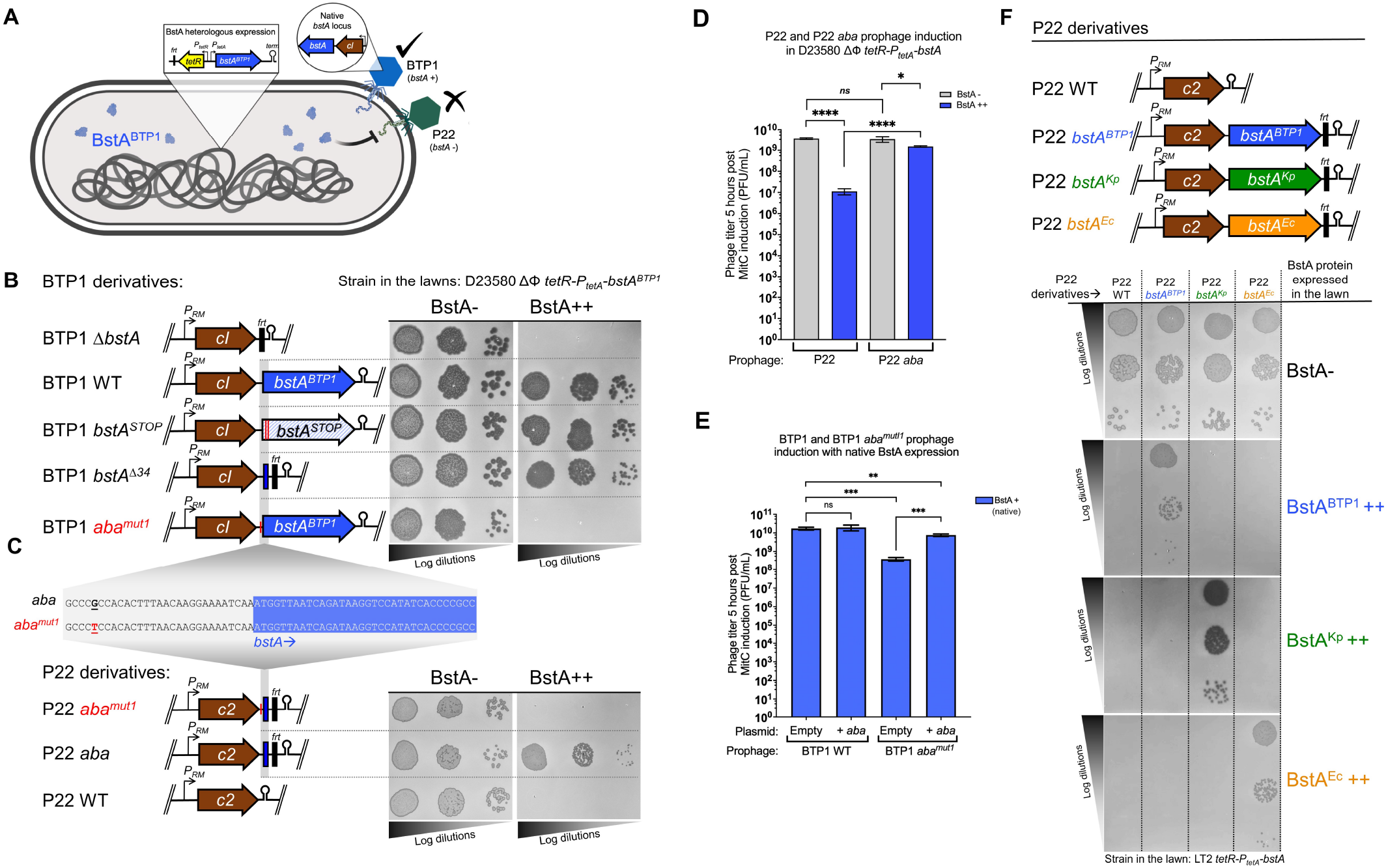
BstA systems include cognate self-immunity elements, *aba*, which are required for successful prophage induction. (**A**) Cartoon summarising the data from Figure 1E. The BTP1 phage, which encodes the *bstA* locus, is not affected by heterologous expression of BstA^BTP1^, whilst the replication of phage P22 is inhibited. (**B**) Schematic of the BTP1-derived phages used and the corresponding effect on sensitivity to BstA^BTP1^ expression (plaque assay). (**C**) Schematic of the P22-derived phages used. In all cases, introduced sequences (*bstA* homologs or fragments) were inserted downstream of the *c2* repressor gene of P22 and are linked to the *frt* sequence. Hairpins represent Rho-independent terminators. Insensitivity of the BTP1 phage to BstA^BTP1^ is dependent on the *bstA* locus on the phage chromosome. However, only the first 34 bp of the *bstA* gene are required, along with 29 bp upstream (in total a 63 bp sequence). The *aba* sequence (native in BTP1, or engineered into P22) counteracts the BstA-driven phage resistance. The GàT mutation (*aba^mu1^*) causes loss of *aba* function, and suppresses the anti-*bstA* interference. (**D**) BstA represses P22 prophage induction in the absence of *aba*. P22 induction was measured in D23580 ΔΦ *tetR-P_tetA_-bstA^BTP1^* lysogenized with prophages P22 WT or P22 *aba* (strain SNW583 and SNW585, respectively). The induced phage titer was measured 5 hours post induction with Mitomycin C (MitC). (**E**) Endogenous BstA represses BTP1 prophage induction in the presence of the *aba^mut1^* mutation, but replication can be rescued by supply of a functional *aba* in *trans*. Prophage induction was measured in strain D23580 δΦ ΔTn*21* (Ap^S^) lysogenized with BTP1 WT (*aba^WT^*) or BTP1 *aba^mut1^* (strain SNW597 and SNW598, respectively). Lysogens were transformed with pUC18 (vector) or pUC18-*aba* (pNAW203, +*aba*) and prophage induction was measured 5 hours post-induction with Mitomycin C. Data in D & E are presented as the mean of biological triplicates ± SD. Groups were compared using unpaired two-tailed Student *t*-test and P values and significance is indicated by ********** or ns (not significant). (**F**) Each *bstA* locus encodes a homolog-specific anti-*bstA* element (*aba*) that suppresses BstA-mediated phage defense. Transfer of each *bstA* locus to phage P22 only confers immunity against the cognate BstA protein. Plaque assays were carried out with the indicated phages, applied on mock-induced (BstA-) or AHT-induced (BstA++) lawns of the indicated strain: SNW576 for panels B & C and strains JH4400 (BstA^BTP1^++), JH4404 (BstA^Kp^++) or JH4408 (BstA^Ec^ ++) for panel F.

To identify the precise genetic basis of BstA self-immunity, we constructed a series of BTP1 mutant phages, carrying truncations of different lengths from the 3’ end of the *bstA* locus (Supplementary Figure 5A) and screened these phages for the ability to replicate in the presence of BstA^BTP1^ expression. Self-immunity (i.e. insensitivity to BstA^BTP1^ expression) was preserved in all mutant phages, except the mutant with the longest *bstA* truncation (BTP1 *bstA^Δ24^*), in which just the first 24 bp of the *bstA* reading frame were intact (Supplementary Figure 5A). A similar truncation mutant containing just the first 34 bp of *bstA* (BTP1 *bstAΔ^34^*) retained immunity to BstA, suggesting that the first 34 bp of the *bstA* gene are essential for the activity of the anti-BstA determinant. Consistently, the transfer of *bstA^Δ34^* (the first 34 bp of *bstA*, along with the upstream sequence) to phage P22 (P22 *bstA^Δ34I^*), conferred BstA immunity (Supplementary Figure 5B). To identify the minimal sequence required for BstA self-immunity, we constructed further P22 *bstA^Δ34I^*-derived phages, successively truncating the transferred sequence from the 5’ end (P22 *bstA^Δ34I^-P22 bstA^Δ34V^*, Supplementary Figure 5B). We discovered that a 63 bp sequence (GCCCGCCACACTTTAACAAGGAAAATCAA<u>ATG</u>GTTAATCAGATAAGGTCCATATCACCCCGCC) spanning 29 bp of the upstream region and the first 34 bp of the *bstA* coding sequence (start codon underlined) was necessary and sufficient to confer the self-immunity (Figure 6C). We designated this 63 bp element *‘aba’*, for <u>a</u>nti-<u>B</u>st<u>A</u>. Supplying the 63 bp *aba* sequence on the high-copy number pUC18 plasmid (pUC18-*aba*) rescued P22 phage replication in the presence of BstA protein, demonstrating that the self-immunity effect of *aba* is retained in *trans* (when *aba* is not carried by the targeted phage, but is supplied on another replicative element) (Supplementary Figure 6A). The intracellular localisation of BstA protein following phage infection was unaffected by the presence of the pUC18-*aba* plasmid (Supplementary Figure 6B).

### The *aba* element is DNA-based

In the native BTP1 prophage, the *aba* sequence overlaps the start of the *bstA* gene, preventing mutational disruption of the *aba* element without modification of the BstA protein sequence. We therefore used the plasmid *trans*-complementation system (wherein the BstA protein and the *aba* sequence are independently encoded) to probe the function of the *aba* sequence (Supplementary Figures 6A). A notable feature of the 63 bp *aba* sequence is the presence of a direct “CCCGCC” repeat at the terminal ends, which we hypothesised was functionally important. Single nucleotide exchange of the CCC<u>G</u>CC→CCC<u>T</u>CC in the first and second repeat (*aba^mut1^* and *aba^mut2^*, respectively) abolished the self-immunity function of the *aba* element, both when located on a phage (Figure 6B) and from a plasmid *in trans* (Supplementary Figure 7A), showing that the *aba* terminal direct repeats are required for *aba* function. Plasmid-borne expression of BstA efficiently supressed plaque formation of P22 and BTP1 phages lacking a functional *aba* sequence (P22 WT, BTP1 *ΔbstA* or BTP1 *aba^mut1^*) but had no effect on BTP1 WT (Supplementary Figure 7B).

The *aba* plasmid *trans*-complementation system additionally allowed us to interrogate the genetic nature of the *aba* element, which we hypothesised was either DNA, RNA or peptide-based. Though three short open reading frames exist within the *aba* sequence, non-synonymous mutation of the reading frames did not ablate *aba* function (Supplementary Figure 8A), suggesting the *aba*-driven immunity is not mediated by a short peptide. Secondly, we tested whether transcription of *aba* was necessary for immunity. The *aba* sequence was cloned in either orientation into the *Salmonella* chromosome, downstream of the arabinose-inducible *P_BAD_* promoter (D23580 ΔΦ *tetR-bstA^BTP1^ P_BAD_-aba*; Supplementary Figure 8B) to produce high levels of *aba* RNA transcripts. A *gfp* gene was inserted downstream to report transcription form the *P_BAD_* promoter. High-level transcription of *aba* RNA did not restore P22 or 9NA plaque formation in the presence of BstA protein, suggesting that transcription of *aba* is not required for anti-BstA activity. This experiment also showed that a single chromosomal copy on *aba* does not confer self-immunity (Supplementary Figure 8B), suggesting that in *trans* (i.e. not encoded on the phage chromosome), *aba* can only suppress BstA when supplied on high-copy replicative elements. Further mutational disruption of the *aba* sequence revealed that the self-immunity function was sensitive to mutation at multiple sites in the 63 bp sequence (Supplementary Figure 8C). Collectively, our data suggest that *aba*-driven suppression of BstA is neither peptide or transcript-mediated, and supports a model where BstA suppression is mediated by *aba* DNA.

### The *aba* element prevents the *bstA*-encoding prophage from aborting its own lytic replication

Unlike most mechanistically-characterised abortive infection systems, a unique feature of the BstA system is its frequent occurrence on prophages (latent forms of active phages) (Figure 2A). Prophages must be able to switch to lytic replication, or else the prophage-state becomes an evolutionary dead-end for the phage. Consequently, prophages which encode defense systems targeting later stages of phage replication (i.e. those that are shared by the induced prophage; DNA-replication onwards) must possess a mechanism of self-immunity. Such self-immunity functions in prophage-encoded phage defense systems have not been previously described.

We hypothesised that the primary biological role of the *aba* element is to allow the endogenous *bstA*-encoding phage to escape BstA-mediated inhibition upon induction from the prophage-state. To test this, we measured the level of induction of prophage P22 in the presence of heterologously-expressed BstA^BTP1^ protein (Figure 6D). In the absence of BstA^BTP1^ expression, the P22 prophage generated a titer of ~4×10^9^ PFU/mL after 5 hours growth with inducing agent (mitomycin C, MitC). However, with BstA^BTP1^ expression, the MitC-induced titer of P22 dropped >300-fold to ~1×10^7^ PFU/mL, showing that BstA inhibited P22 phage replication. Transfer of the *aba* sequence to prophage P22 (P22 *aba*) significantly increased the induced titer in the presence of BstA^BTP1^ to ~1.5×10^9^, restoring it to the level seen in the absence of BstA, and showing that the *aba* element rescues prophage induction *via* suppression of BstA.

Finally, we validated the importance of the *aba* element in the context of native BTP1 prophage induction. The presence of additional copies of the *aba* sequence *in trans* on the pUC18 plasmid did not affect the titer of BTP1 phage generated after 5 hours growth with inducing agent (MitC), suggesting that native levels of BstA protein do not constrain BTP1 prophage induction in the presence of the native, functional *aba* element (Figure 6E). However, when the *aba^mut1^* mutation (exchange of a single functionally important nucleotide in the terminal direct repeat) was introduced into the BTP1 prophage, the MitC-induced titer of phage BTP1 was reduced ~40-fold in the presence of native BstA expression. This reduction was almost entirely rescued when the *aba* sequence was supplied *in trans* on the pUC18 plasmid, confirming that the *aba^mut1^* mutation ablates the function of the *aba* element. When the *aba^mut1^* mutation was introduced into the BTP1 prophage in the absence of native BstA protein expression (D23580 ΔΦ [BTP1 *aba^mut1^ bstA^STOP^*]) there was no effect on prophage induction (Supplementary Figure 9), confirming that the effect of the *aba^mut1^* mutation is dependent on the presence of BstA.

These intricate experiments demonstrate that a functional *aba* element is required for the *bstA*-encoding prophage to switch from a lysogenic to lytic lifestyle. In the absence of *aba*, the *bstA*-encoding prophage suffers replication inhibition by endogenous BstA protein (self-targeting), presumably by the same abortive infection mechanism that inhibits exogenous phage infection.

### Distinct BstA proteins are associated with cognate *aba* elements

Finally, we determined whether the *aba* sequence from *bstA^BTP1^* could suppress the activity of the BstA proteins of other bacteria. We challenged the P22 *bstA^BTP1^* phage (immune to expression of BstA^BTP1^ due to the presence of *aba*) against expression of BstA^Ec^ or BstA^Kp^. The *bstA^BTP1^* locus did not protect P22 from the heterologous BstA proteins, suggesting that the *aba* element from *bstA*^BTP1^ only confers immunity again BstA^BTP1^, raising the possibility that heterologous BstA proteins have cognate *aba* elements (Figure 6F). To test this hypothesis, we engineered P22 phages to encode either *bstA^Ec^* or *bstA^Kp^* (including the respective upstream sequence). Consistent with a cognate BstA-*aba* interaction, P22 *bstA^Ec^* became specifically immune to expression of BstA^Ec^, and P22 *bstA^Kp^* gained specific immunity to BstA^Kp^ expression (Figure 6F). Each *bstA* locus therefore encodes highly-specific self-immunity. We conclude that whilst BstA proteins are broadly functionally interchangeable in terms of their phage-defense activity, each *bstA* locus contains a cognate *aba* element that is inactive against heterologous BstA proteins. The specificity of the *aba* self-immunity element means that heterologous *bstA*-encoding phages are unable to bypass BstA-mediated abortive infection, making *aba*-mediated suppression of BstA exclusive to the induced *bstA*-encoding prophage.

### BstA protein does not affect phage lysogenic development but inhibits DNA replication during lytic development

To interrogate how the BstA protein interacts with infecting bacteriophages, we determined whether lysogenic phage development, where the infecting phage integrates into the genome of the bacterium, was affected by BstA expression. We previously showed that plaque formation (which is indicative of lytic development) of the temperate phage P22 WT is inhibited by BstA protein (Figure 1E). We used an antibiotic-tagged derivative of P22 (P22 *Δpid::aph*) to determine the frequency of lysogeny with and without BstA expression. We found that the frequency of lysogeny was approximately 6% (Figure 7A) regardless of the presence of BstA, suggesting that BstA expression does not affect phage lysogenic development. This finding suggests that BstA activity is triggered by, or targets, an aspect of phage lytic replication not shared by lysogenic development. Further, it implies that BstA has no effect on initial stages of phages infection that occur prior to lysogenic development i.e. adsorption and DNA translocation.

**Figure 7:**
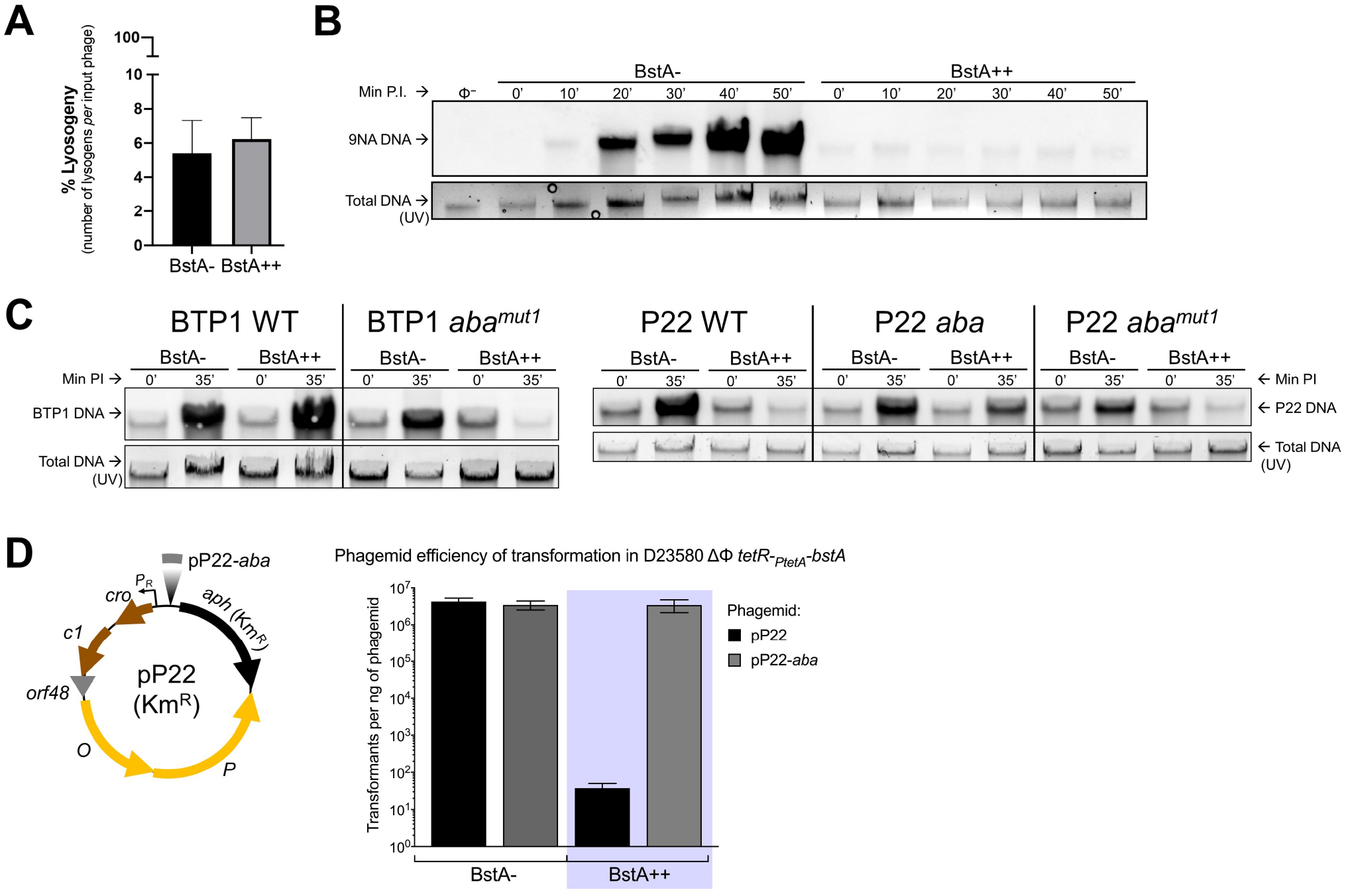
BstA protein does not affect phage lysogeny but inhibits phage DNA replication in the absence of *aba*. (**A**) Frequency of lysogeny of the P22 *Δpid::aph* phage in mock-induced (BstA-) or AHT-induced (BstA++) D23580 ΔΦ *tetR-P_tetA_-bstA^BTP1^* (SNW576). DNA replication of 9NA phage (**B**), BTP1 and P22-derived phages (**C**) in the absence or presence of BstA expression. Phage DNA was detected by Southern blotting with total DNA extracted from mock-induced (BstA-) or AHT-induced (BstA++) host strain D23580 ΔΦ *tetR-P_tetA_-bstA^BTP1^* (SNW576), infected by the indicated phage at MOI=5. Before the transfer procedures, total stained DNA was visualized from gels under UV light and the resulting pictures served as loading control. Min PI = minutes post infection. Non-infected SNW576 DNA was used as negative control to check the DNA probe specificity. (**D**) *aba* dramatically increases the transformation efficiency of P22 derived phagemids in BstA expressing *Salmonella.* The Km^R^ phagemids pP22 (pNAW229) and pP22-aba (pNAW230) are schematized, and the efficiency of transformation for each phagemid was measured in mock-or AHT-induced competent bacteria of strain D23580 ΔΦ *tetR-P_tetA_-bstA^BTP1^* (SNW576). Data are presented as the mean of biological triplicates ± SD.

Sequence-based analysis of BstA protein homologs suggested that the N-terminal domain may bind DNA (Figure 2C), and fluorescence microscopy showed BstA protein co-localizing with phage DNA (Figure 5C). The replication of DNA is crucial for phage morphogenesis, as a new copy of the phage chromosome is required for packaging into the capsid of each new virion. To test whether BstA protein inhibits phage DNA replication during lytic development in a manner that can be suppressed by *aba*, we conducted Southern blot experiments to monitor levels of phage DNA during infection. During successful phage replication, phage DNA accumulates in the cell. Using our prophage-negative, inducible BstA-expression strain (D23580 ΔΦ *tetR-P_tetA_-bstA^BTP1^*), we first tested the replication of the BstA-sensitive virulent phage, 9NA. In the absence of BstA expression, the level of phage 9NA DNA gradually increased over a 50 minute infection time course, reflecting successful phage replication (Figure 7B). However, no accumulation of phage 9NA DNA was observed in the presence of BstA^BTP1^, suggesting that BstA protein strongly inhibited the replication of phage DNA.

Consistent with the self-immunity function of *aba*, BTP1 phage DNA replication was not affected by the expression of BstA^BTP1^, unless the *aba* element was non-functional (BTP1 *aba^mut1^*) (Figure 7C). Likewise, successful replication of phage P22 DNA in the presence of BstA^BTP1^ only occurred when the phage possessed a functional *aba* element (Figure 6B).

To confirm that BstA protein inhibits DNA replication, we constructed small phage-derived plasmids (‘phagemids’) based on the phage P22 replication module (pP22) (Figure 7D), and a P22 phagemid that included the 63 bp *aba* sequence (pP22-*aba*).

*Salmonella* cells were transformed with the phagemids in the presence or absence of BstA^BTP1^ protein expression. In the absence of BstA, the stable replication of both P22 phagemids in *Salmonella* cells generated >10^6^ transformants/ng phagemid. However, expression of BstA^BTP1^ reduced the transformation efficiency of pP22 (lacking the *aba* sequence) to around 10 transformants/ng. Addition of the *aba* sequence to the phagemid (pP22-*aba*) restored the transformation efficiency of the phagemid in the presence of BstA to BstA-negative levels (Figure 7D).

We conclude that phage DNA replication is strongly suppressed by BstA, but replication can be rescued by the *aba* element, presumably by suppression of BstA protein activity. As replicated phage DNA is an essential substrate for packaging into phage capsids, the inhibition of DNA replication is likely to prevent the production of infectious progeny phages, consistent with the observation that infectious phages are not released from BstA-expressing cells following cell lysis (Figure 4). We propose that the activity of BstA protein supresses phage DNA replication, a process that can be blocked by the native prophage carrying the *aba* self-immunity element.

## Discussion

Prophages are latent phages residing in the genomes of and frequently encode accessory genes that bestow beneficial functions on their host bacteria (Bondy-Denomy and Davidson, 2014; Cumby et al., 2012). A function that can significantly increase bacterial fitness in many environments is phage resistance, and prophages may be a large reservoir of uncharacterised phage defense systems (Dedrick et al., 2017; Snyder, 1995; Tsao et al., 2018).

Here, we have discovered a novel family of prophage-encoded abortive infection proteins (BstA) which efficiently defend bacterial populations from phage epidemics. BstA protein is constitutively expressed inside cells that carry the prophage, and provides effective population-level phage defense through abortive infection, inhibiting phage replication at the cost of the viability of individual infected cells. Possession of such innate phage defense systems by active prophages imposes an obvious challenge: the prophage must avoid self-targeting by its own defense system when switching to lytic replication.

We realised that the native prophage which encodes BstA required a mechanism to counteract the protein upon induction from the prophage-state, to avoid aborting its own lytic replication. The BstA system solves this problem with the *aba* element (<u>a</u>nti-<u>B</u>st<u>A</u>), a co-encoded short DNA sequence that specifically suppresses the activity of BstA protein upon prophage induction, giving the induced prophage self-immunity against endogenous BstA protein. Theoretically, such a system might leave BstA-expressing cells vulnerable to infection by heterologous BstA-encoding phages, which could use their own *aba* element to bypass native BstA. This problem is avoided by cognate BstA-*aba* pairs, as each BstA protein is suppressed only by the cognate *aba* element, ensuring that BstA supression is specific to the native BstA-encoding prophage.

Despite over 60 years of investigation, abortive infection systems remain mysterious, and very few have been characterised more deeply than the level of broad functional mechanism (Labrie et al., 2010). Here, we present a high-level overview of the BstA phage defense system, and the corresponding anti-BstA *aba* element. We are left with two major questions regarding the activity of the BstA protein. Firstly, what are the phage determinants for BstA sensitivity? Though BstA was active against approximately 50% of the phages tested, we did not detect similarities between BstA-targeted and non-targeted phages that could reflect the molecular determinants of sensitivity. It remains possible that rather than responding to a physical phage stimulus, such as phage DNA or protein, BstA protein responds to a cellular stimulus produced by the infection of specific types of phages, for example the recruitment of DNA replication machinery.

Secondly, what is the molecular mechanism by which BstA protein inhibits phage DNA replication? Our data suggest that phage DNA does not replicate in the presence of BstA. Though numerous Abi systems in *Lactococcus* have been proposed to interfere with phage DNA replicative functions (Chopin et al., 2005), the molecular mechanisms have not been well characterised. The existence of a putative DNA-binding domain in BstA proteins, and microscopic observation of BstA co-localisation with phage DNA make it tempting to speculate that BstA interacts physically with phage DNA to prevent replication, for example by occlusion of a replication initiation site.

Alongside the mechanistic details of the BstA protein that remain to be established, little is known about the interaction of BstA with the *aba* element. Our data show that *aba* interacts with BstA in DNA form, but the mechanism by which *aba* DNA suppresses BstA protein is unclear. Our data indicate that multiple copies of the *aba* element are required to suppress BstA protein in *trans*, which could explain why *aba* function is specific to prophage induction (when prophage DNA is replicated to a high copy number). However, copy-number cannot be the only factor affecting *aba* functionality, because an induced prophage is evidently able to suppress BstA protein right from the beginning of prophage induction, when *aba* is present as just a single copy on the chromosome. Further study of the BstA-*aba* system is required to resolve the precise molecular mechanisms by which BstA-encoding prophages, such as BTP1, achieve self-immunity.

We consistently observed that phage-infected cells that contained functional BstA protein underwent lysis, probably in the absence of infectious progeny phage release. However we cannot be certain whether BstA protein acts actively or passively to cause cell lysis. Abi systems have frequently been termed “altruistic suicide” systems, which mediate “programmed cell death” in response to phage infection (Abedon, 2012; Shub, 1994). Whilst perhaps a useful conceptual analogy for the strictly population-level effect of Abi systems, this narrative implies that Abi systems actively cause cell death. Though this is likely to sometimes be the case, such as in the CBASS system (Cohen et al., 2019), Abi can also be achieved by simple disruption of the phage replication pathway. Because phage lysis is generally a temporally programmed event that occurs independently of successful virion morphogenesis (Cahill & Young, 2018), phage-mediated cell lysis can occur in the absence of virion assembly. For example many *Lactococcus* Abi systems target aspects of phage replication, such as AbiZ which is thought to interact with phage holin proteins, to stimulate premature cell-lysis before virion assembly is completed (Durmaz and Klaenhammer, 2007).

It is possible that BstA protein simply inhibits viable phage particle formation, for example by precluding the formation of concatemeric DNA for packaging, whilst allowing the phage lytic pathway to proceed unperturbed to cell lysis. However, inhibition of phage DNA replication would dramatically reduce substrates for transcription and translation of phage lysis gene products, yet we did not observe a difference in the timing of cell lysis for phage infected cells in the presence or absence of BstA during microscopy studies. The exact mechanism of cell lysis during BstA Abi activity will require further study.

An intriguing feature of the BstA phage defense system is its tight association with prophages, and specifically, with the prophage repressor locus. Though we found homologs in diverse Gram-negative bacteria, the genetic architecture of the *bstA* locus (i.e. lying downstream of, and presumably sharing the promoter of the prophage repressor) was strikingly conserved. The region between the repressor (*cI*) and *n* gene of lambdoid phages has previously been identified as a hotspot of mosaic diversity (Degnan et al., 2007). In fact, the corresponding site in phage Lambda harbours the *rexAB* genes, perhaps the most widely studied prophage-encoded abortive infection system (Snyder, 1995). Despite >60 years of research, the molecular mechanisms of RexAB activity are poorly understood. RexB is reported to be an ion channel, which triggers loss of cell membrane potential upon activation by the intracellular sensor RexA (Labrie et al., 2010; Snyder, 1995). Whilst not mechanistically comparable to BstA, perhaps the shared synteny of the BstA and RexAB abortive infection systems points to a functional significance of this genomic region, as the *cI* repressor gene is one of the most highly transcribed prophage promoters during lysogeny.

Though somewhat functionally analogous to toxin-antitoxin systems, to our knowledge no other example of self-immunity mechanisms have been described within prophage-encoded abortive infection systems. However, some evidence supports the widespread existence of such mechanisms. For example, the activity of Lambda RexB protein can be suppressed by overexpression of the *rexB* gene relative to *rexA*. It has been speculated, but not shown experimentally, that high levels of RexB might allow phage Lambda to replicate lytically in the presence of RexAB (Parma et al., 1992) i.e. giving the Lambda prophage self-immunity against its own Abi proteins. Our discovery of the BstA and the *aba* element supports this hypothesis.

In conclusion, the discovery of the BstA-*aba* system opens unexplored avenues of research into the mechanisms used by prophages to suppress their own phage-defense activities. We anticipate that similar strategies may be widespread and commonplace, perhaps existing within prophage-encoded phage defense systems that have already been identified. Given the huge mosaic diversity of temperate phages, and high prevalence of uncharacterised accessory genes, the reservoir of prophage-encoded phage defense and self-immunity systems is likely to be vast and largely unexplored.

## Methods

### RESOURCE AVAILABILITY

Further information and requests for bacterial and bacteriophage strains should be addressed to the corresponding author.

### EXPERIMENTAL MODEL AND SUBJECT DETAILS

#### Bacteria and bacteriophages

The full list of bacterial strains used and constructed is available in Supplementary Table 2. All the *Salmonella* strains were derived from the African *S.* Typhimurium ST313 strain D23580 (GenBank: FN424405.1) (Kingsley et al., 2009) or the model *S.* Typhimurium strain LT2 (GenBank: AE006468.2)(McClelland et al., 2001b; Zinder and Lederberg, 1952). All the *Escherichia coli* strains constructed were derived from *E. coli* strain K-12 substrain MG1655 (GenBank: NC_000913.3) (Riley et al., 2006). The *bstA* homolog genes were cloned from *E. coli* NCTC10963 (GenBank: NZ_CAADJH010000002.1) or from *K. pneumoniae* Kp52.145 (GenBank: FO834906.1) (Bialek-Davenet et al., 2014). Bacteriophages (phages), including the temperate phages P22 (GenBank: NC_002371.2) (Pedulla et al., 2003) and BTP1 (GenBank; NC_042346.1) (Owen et al., 2017) and their derivatives, are described in Supplementary Table 2. The genomic coordinates and gene identifiers indicated below refer to the GenBank accession numbers mentioned above.

## METHOD DETAILS

### Growth conditions and transformation

All suppliers of chemical and reagents are specified in the Key Resources Table. Unless stated otherwise, bacteria were grown at 37°C in autoclaved Lennox Broth (LB: 10 g/L Bacto Tryptone, 5 g/L Bacto Yeast Extract, 5 g/L NaCl) with aeration (shaking 220 rpm) or on LB agar plates, solidified with 1.5% Agar. The salt-free LBO media contained 10 g/L Bacto Tryptone, 5 g/L Bacto Yeast Extract. Pre-cultures were inoculated with isolated colonies from agar plates and grown to stationary phase (for at least 6 hours) in 5 mL LB in 30 mL universal glass tubes or in 50 mL plastic tubes (Greiner).

Cultures were typically prepared by diluting the pre-cultures (1:100) or (1:1000) in LB, and bacteria were grown in conical flasks containing 10% of their capacity of medium (*i.e.* 25 mL LB in a 250 mL conical flask) with aeration. For fluorescent microscopy experiments, bacteria were grown in M9 minimal medium (Sambrook and Russell, 2001) supplemented with 0.4% glucose and 0.1% Bacto Casamino Acids Technical (M9 Glu^+^).

When required, antibiotics were added to the media: 50 μg/mL kanamycin monosulfate (Km), 100 μg/mL Ampicillin sodium (Ap), 25 μg/mL tetracycline hydrochloride (Tc), 20 μg/mL gentamicin sulfate (Gm), 20 μg/mL chloramphenicol (Cm). Bacteria carrying inducible constructs with genes under the control of the *P_BAD_* or *P_m_* promoters were induced by adding 0.2 % (w/v) L-(+)-arabinose or 1 mM *m*-toluate, respectively. For the strains carrying *tetR-P_tetA_* modules, *P_tetA_* induction was triggered by adding 500 ng/mL of anhydrotetracycline hydrochloride (AHT, stock solubilized in methanol). For these constructs, the same volume of methanol was added to the non-induced cultures (mock treatment). Chemically-competent *E. coli* were prepared with RbCl-based solutions and were transformed by heat shock (Green and Rogers, 2014).

For the preparation of electro-competent cells, bacteria were grown in the salt-free medium LBO to an Optical Density at 600 nm (OD_600_) of 0.4-0.5. The bacteria were washed twice with cold sterile Milli-Q water (same volume as the culture volume) and were concentrated 100 times in cold 10% glycerol, prior to storage at −80°C. When ultra-competent *Salmonella* cells were required, the bacteria were grown in LBO at 45°C to OD_600_ 0.4-0.5, because growth at high temperature inactivates the *Salmonella* restriction systems (Edwards et al., 1999). Competent cells (10-50 μL) were mixed with 10-5000 ng of DNA in electroporation cuvettes (2 mm gap) and the reactions were electroporated (2.5 kV) using a MicroPulser electroporator (Bio-Rad). Bacteria were re-suspended in 0.5-1 mL LB and incubated for recovery at 37°C (30°C for temperature sensitive plasmids) with aeration, for at least one hour. Finally, the transformed bacteria were spread on selective LB agar plates and transformant colonies were obtained after at least 12 hours incubation at 30-37°C.

For assessment of strain growth kinetics with BstA expression, a FLUOstar Omega plate reader (BMG LABTECH) was used as follows: bacteria were inoculated at an initial OD_600_ of 0.01 (six replicates) in 200 μL of LB or LB + AHT in 96-well plates (Greiner). Bacteria were grown at 37°C with aeration (500 rpm, orbital shaking) and the OD_600_ was monitored every 15 min for 15 hours. Uninoculated LB medium was used as blank.

### Cloning procedures

All the plasmids and DNA oligonucleotides (primers) are listed in Supplementary Table 2. DNA manipulation and cloning procedures were carried out according to the enzyme and kit supplier recommendations and to standard procedures (Sambrook and Russell, 2001). DNA purity and concentration were measured with a DeNovix DS-11 FX spectrophotometer/fluorometer and using the Qubit dsDNA HS assay Kit.

For all the cloning procedures, Polymerase Chain Reactions (PCRs) were performed with the Phusion High Fidelity DNA polymerase, purified template DNA and primers in the presence of 3 % Dimethyl Sulfoxide and 1 M betaine, when required. Prior to Sanger sequencing of the constructs, PCR reactions were carried out directly from bacteria or phages with MyTaq Red Mix 2X. PCR fragments were analysed by electrophoresis, purified and finally sequenced with the appropriate primers (Lightrun service, Eurofins Genomics) (Supplementary Table 2).

All the plasmids were constructed as detailed in the Supplementary Table 2 and were verified by Sanger sequencing. Insertions of DNA fragments into plasmids were performed by digestion/ligation procedures, using restrictions enzymes and the T4 DNA ligase. In addition, PCR-driven restriction-free cloning techniques were used: overlap extension PCRs (Heckman and Pease, 2007) and plasmid assembly by PCR cloning (Van Den Ent and Löwe, 2006) were performed with chimeric primers, purified template DNA and Phusion DNA polymerase, as described previously (Owen et al., 2020). Cloning reactions were transformed by heat shock into *E. coli* Top10 (Invitrogen) or S17-1 *λpir* (Simon et al., 1983). New template plasmids were constructed to insert fluorescent protein encoding genes into *Salmonella* or *E. coli* chromosomes, as reported previously (Gerlach et al., 2007). These plasmids carry the *oriR6K* γ origin of replication of pEMG, the *frt-aph-frt* (Km^R^) module of pKD4 linked to *gfp^+^* (pNAW52), *_sf_gfp* (encoding for superfolder GFP, pNAW62) or *mcherry* (pNAW73), amplified respectively from plasmids pZEP09 (Hautefort et al., 2003), pXG10-SF (Corcoran et al., 2012) and pFCcGi (Figueira et al., 2013). A similar template plasmid, carrying the *frt-aph-frt-tetR-P_tetA_* module (pNAW55) was constructed and was used to insert the *tetR* repressor and the AHT-inducible promoter *P_tetA_* upstream of genes of interest, as reported earlier (Schulte et al., 2019). For the construction of gentamicin resistant plasmids, the *aacC1* resistance gene was obtained from plasmid pME4510 (Rist and Kertesz, 1998).

The high copy number plasmid pUC18 was used to clone the different versions of the anti-*bstA* (*aba*) fragment: the *aba* fragments (*aba1-aba14* alleles) were amplified by PCR, digested with EcoRI and BamHI and ligated into the corresponding sites of pUC18. Phagemids based on the phage P22 replication module were constructed by EcoRI/KpnI digestion and ligation, as follows: the *PR* promoter and the *cro-c1-orf48-O-P* genes of P22 (coordinates 31648-34683) were amplified and circularized by ligation with the *aph* Km^R^ cassette of pKD4 or with the *aba-aph* modules, amplified from strain SNW617. The ligations reactions were purified and electroporated into ultra-competent SNW555, a prophage-free and plasmid-free derivative of *S.* Typhimurium D23580. The resulting phagemids pNAW229 (pP22-*aph*), pNAW230 (pP22-*aba*-*aph*) were obtained after selection on Km medium.

Phage DNA was extracted from high titer lysates in LBO: nine volume of the phage lysates were mixed with one volume of 10 X DNase buffer (100 mM Tris-HCl, 25 mM MgCl_2_, pH 7.5) supplemented with RNase A (40 μg/mL final) and DNase I (400 μg/mL final). After 1 hour incubation at 37°C, DNase I was heat-inactivated at 75°C for 10 min and phage DNA was extracted from 500 μL of the nuclease-treated lysates with the Norgen Phage DNA Isolation after Proteinase K treatment, as specified by the manufacturer.

### Genome editing techniques

Strain constructions are detailed in Supplementary Table 2. For chromosomal insertions and deletions, *λ red* recombination was carried out with the arabinose-inducible plasmid pKD46 (for *E. coli)* or with the heat inducible plasmid pSIM5-*tet* (for *Salmonella*), both expressing the *λ red* genes. Bacteria were grown to exponential phase in LBO, according to the resistance and induction condition of the respective *λ red* plasmid (Datsenko and Wanner, 2000; Hammarlöf et al., 2018; Koskiniemi et al., 2011) and electro-competent cells were prepared as mentioned above. PCR fragments carrying a resistance cassette were constructed by overlap extension PCR or were directly obtained by PCR from the appropriate plasmid or strain. Electro-competent cells (40-50 μL) were transformed with 500-5000 ng of the PCR fragments and the recombinants were selected on selective LB agar plates.

Mutations or insertions linked to selective markers were transduced into *Salmonella* strains using the P22 HT *105/1 int-201* (P22 HT) transducing phage (Owen et al., 2017; Schmieger, 1972). For *E. coli*, the transducing phage P1 *vir* was used (Ikeda and Tomizawa, 1965; Tiruvadi Krishnan et al., 2015). Transductants were grown on selective LB agar plates supplemented with 10 mM EGTA. After two passages, clearance of the transducing phages was confirmed by diagnostic PCR using primer pairs NW_62/NW_63 for P22 HT or NW_392/NW_393 for P1 *vir* and by a passage on Green Agar medium (Maloy, 1990).To remove the antibiotic cassettes, flanked by FLP recognition target sites (*frt*), the FLP recombinase expressing plasmids pCP20, pCP20-TcR and pCP20-Gm were used, as previously reported (Cherepanov and Wackernagel, 1995; Doublet et al., 2008; Hammarlöf et al., 2018; Kintz et al., 2015). The inducible *tetR-P_tetA_-bstA* modules were constructed by fusing the *frt-aph-frt-tetR-P_tetA_* module of pNAW55 to the *bstA* gene of D23580 (*bstA^BTP1^, STMMW_03531*), *E. coli* NCTC10963 (*bstA^Ec^, E4V89_RS07420*) or *K. pneumoniae* Kp52.145 (*bstA^Kp^, BN49_1470*). Each construct carries the native *bstA* ribosome binding site and Rho-independent terminator. The *tetR-P_tetA_-bstA* modules were inserted by *λ red* recombination into the *STM1553* pseudogene of *S.* Typhimurium LT2 (between coordinates 1629109-1629311), corresponding to *STMMW_15481* in D23580 (coordinates 1621832-3). Previously we have shown that the *STM1553* and *STMMW_15481* genes are not expressed at the transcriptional level (Canals et al., 2019).

In *E. coli* MG1655, the *bstA* modules were inserted into the *glmS-pstS* intergenic region (coordinates 3911773-4). To generate Ap and Cm sensitive D23580 strains, the pSLT-BT plasmid-encoded Tn*21*-like element, that carries the resistance genes (Kingsley et al., 2009), was replaced by the Km^R^ cassette of pDK4 by *λ red* recombination (deletion coordinates 34307 to 57061, GenBank: NC_013437.1). The resulting large single-copy plasmid pSLT-BT *ΔTn21::aph* was extracted (Heringa et al., 2007) and electroporated into the strains of interest. After selection on Km medium, the Ap and Cm sensitivity was confirmed and the Km^R^ cassette was flipped out using pCP20-Gm. For scarless genome editing, the pEMG plasmid-based allelic exchange system was used (Martínez-García and de Lorenzo, 2011). The pEMG derivative suicide plasmids were constructed as specified in Supplementary Table 2 and were replicated in *E. coli* S17-1 *λpir*. Conjugation of the resulting plasmids into *Salmonella* and subsequent merodiploid resolution with plasmid pSW-2 were carried out as previously described (Canals et al., 2019; Owen et al., 2017). Some key strains and phages (indicated in Supplementary Table 2) used in this study were whole-genome sequenced (Illumina) at MicrobesNG (Birmingham, UK).

### Plasmid deletion in *S.* Typhimurium D23580

The pSLT-BT, pBT1, pBT2 and pBT3 plasmids (Kingsley et al., 2009) were cured from strain D23580, using the CRISPR-Cas9-based methodology (Lauritsen et al., 2017). A novel CRISPR-Cas9 Km resistant plasmid (pNAW136) was obtained by ligating the CRISPR-Cas9 module of plasmid pCas9 (Jiang et al., 2013) with the unstable origin of replication *ori*RK2, the *trfA* replication gene and the *aph* Km^R^ gene. Anti-plasmid protospacers (30 bp) were generated by the annealing of 5’-phosphorylated primer pairs that targeted the pSLT-BT, pBT1, pBT2 and pBT3 plasmids, designed according to the Marraffini Lab protocol (Jiang et al., 2013). The protospacers were ligated into *Bsa*I-digested pNAW136 with T4 DNA ligase and the resulting plasmids were checked by Sanger sequencing, using primer NW_658.

The resulting plasmids pNAW168 (anti-pSLT-BT) and pNAW169 (anti-pBT1), pNAW139 (anti-pBT2) and pNAW191 (anti-pBT3) were electroporated into D23580-derived strains and transformants were selected on Km plates. After two passages on Km, the loss of the pSLT-BT, pBT1, pBT2 or pBT3 plasmids was confirmed by diagnostic PCR. The absence of the unstable pNAW136-derived plasmids was confirmed by the Km sensitive phenotype of colonies after two passages on non-selective medium.

### Phage stock preparation and plaque assays

All phage stocks were prepared in LB or LBO. For *Salmonella* phages, the prophage-free strain *S.* Typhimurium D23580 ΔΦ (JH3949) was used as host (Owen et al., 2017). Exponential phase cultures of D23580 ΔΦ were infected with ~10^5^ Plaque Forming Units (PFU) and infected cultures were incubated for at least 3 hours at 37°C (with aeration). Phages lysates were spun down (4,000 X *g*, 15 min) and supernatants were filter-sterilized (0.22 μm, StarLab syringe filters). The resulting phage lysates were stored at 4°C in the presence 1% chloroform to prevent bacterial contamination.

Coliphage lysates were prepared similarly with *E. coli* MG1655 as host. When required, maltose (0.2%), CaCl_2_ (10 mM) and MgSO_4_ (10 mM) were added during the infection (λ, P1 *vir* and Φ80*p*SU3^+^). For Φ80-derived phages, the infection temperature was reduced to 30°C (Rotman et al., 2010).

Phage lysates were serial-diluted (decimal dilutions) with LB and virion enumeration was performed by double-layer overlay plaque assay (Kropinski et al., 2009), as follows. Bacterial lawns were prepared with stationary phase cultures of the reporter strains, diluted 40 times with warm Top Agar (0.5 % agar in LB, 50°C). The seeded Top Agar was poured on LB 1.5% agar bottom layer: 4 mL for 8.6 cm diameter petri dishes or 8 mL for 12 x 12 cm square plates.

When inducible *P_tetA_* or *P_BAD_* constructs were present in the reporter bacteria, 500 ng/mL of AHT or 0.2 % arabinose were added in the Top Agar. When required, antibiotics were added in the Top Agar layer. The bacterial lawns were incubated for 30 min at room temperature with the appropriate inducer, to allow solidification and the expression of the inducible genes. Finally, phages suspensions (5-20 μL) were applied on the Top Agar surface and pictures of the resulting plaques were taken with an ImageQuant Las 4000 imager (GE Healthcare) after 16-20 hours incubation at 30 or 37°C.

### Construction of P22 virulent phages

For the generation of obligately virulent P22 phages, a 633 bp in-frame deletion (coordinates 31028-31660) was introduced in the *c2* repressor gene by *λ red* recombination in a P22 lysogen as follows. Two fragments of ~500 bp, flanking *c2*, were amplified with primers pairs NW_818 / NW_819 and NW_820 / NW_821. The two amplicons were fused by overlap extension PCR and 1000-3000 ng of the resulting *Δc2* fragment were electroporated into P22 lysogens (in the prophage-free D23580 ΔΦ background) carrying the *λ red* recombination plasmid pSIM5-*tet*, as described above. The transformation reactions were re-suspended in 5 mL LB and incubated for 2 hours at 37°C with aeration. The culture supernatants were filter sterilized and serial-diluted to 10^-2^. Ten microliters of each dilution were mixed with 100 μL of a D23580 ΔΦ stationary phase culture and with 4 mL of warm Top Agar. The mixtures were poured on LB agar plates and the plates were incubated for ~16 hours at 37°C. P22 Δ*c2* recombinants were identified by the clear morphology of their plaques, compared to the turbid plaques of WT P22. The Δ*c2* deletion was confirmed by PCR and Sanger sequencing with primers NW_406 and NW_805.

### Use of the *Δtsp-gtrAC* genetic background

Where possible, experiments were carried out with native BstA expression (from its natural locus within the BTP1 prophage), to best recapitulate the natural biological activity of the protein. However, as the *gtr* locus of phage BTP1 blocks attachment of many phages including P22 and BTP1, to achieve efficient phage infections we consistently used a strain background where the *gtr* locus has been inactivated (Δ*tsp-gtrAC*). The BTP1 prophage spontaneously induces to a titer of ~10^9^ PFU/mL in liquid culture (Owen et al., 2017), and in the absence of *gtr* activity in surrounding cells, free BTP1 phages mediate cleavage of the O-antigen *via* the putative enzymatic activity of the tailspike protein (Kintz et al., 2015).

Consequently, to avoid an unnatural, short LPS phenotype as a result of *gtr* inactivation in a BTP1 lysogen, we additionally inactivated the upstream gene encoding for the BTP1 tailspike (*tsp*) (D23580 *Δtsp-gtrAC*, JH4287). Full details of the construction of this strain can be found in the Supplementary Table 2.

### Phage replication assay

Stationary phase cultures of the reporter bacteria were diluted to OD_600_ 0.4 with LB. Aliquots (0.2 mL) were prepared in 1.5 mL tubes and phage stock suspensions were added to a final phage titer of 100-1000 PFU/mL. The infections were carried out at 37°C (30°C for Φ80*p*SU3^+^) with shaking for 2-4 hours and were stopped by the addition of 20 μL of chloroform. After a 10 sec vortex, the lysates were centrifuged (20,000 X *g*, 5 min) and serial diluted. When M9 Glu^+^ was used, *Salmonella* strains were grown to OD_600_ ~ 0.5 in this medium prior to phage infection.

Phage titer was determined by plaque assay: 10 μL of the dilutions were applied to bacterial lawns of the appropriate reporter strain in technical triplicates. Plaques were enumerated after 16-20 hours of incubation and phage titers (PFU/mL) were calculated for each lysate. To measure the phage input at time 0 (T^0^), the same volume of stock phage suspension was added to 0.2 mL of bacteria-free LB and the titer was determined as described above. The fold-replication for each phage was calculated as the phage titer of the lysate post infection divided by the input phage titer at T^0^. When the phage titer in the lysate was lower than the phage input, the replication was considered to be null (“<1-fold). When AHT inducible *tetR-P_tetA_-bstA* strains were used, AHT (500 ng/mL) or methanol (mock) were added to the diluted bacterial suspension and phages were added after 15 min of incubation at 37°C with aeration.

For replication assays of the coliphages *λ*, P1 *vir* and Φ80*p*SU3^+^, *E. coli* strains were grown to exponential phase (OD_600_ 0.4) in LB and phages were added as mentioned above. To stimulate infection by these phages, maltose (0.2%), CaCl_2_ (10 mM) and MgSO_4_ (10 mM) were added during the infection and in the lawns of the reporter *E. coli* MG1655. All the phage replication experiments presented were carried out at least twice with biological triplicates.

### Induction of P22 and BTP1 prophages

D23580 ΔΦ-derived lysogens that carried the different versions of P22 and BTP1 were constructed as detailed in the Supplementary Table 2. For complementation with the pUC18-derived plasmids (Ap^R^), Ap sensitive lysogens were constructed by the inactivation of the Tn*21*-like element, as described above. The resulting lysogens were grown to stationary phase in LB and the pre-cultures were diluted 100-1000 times in fresh LB and grown to OD_600_ 0.4-0.5, prior addition of Mitomycin C (MitC, 2 μg/mL). The induced cultures were incubated for 3-5 hours at 37°C with aeration and cultures were filter sterilized and serial diluted. The phage titer was measured by plaque assay on the appropriate host strain lawn with technical replicates, as described above. All the prophage induction experiments were carried out at least twice with biological triplicates.

### Survival assays

For the survival assays, D23580 *Δtsp-gtrAC* (JH4287), D23580 *Δtsp-gtrAC bstA^STOP^* (SNW431) or D23580 ΔΦ [P22] (SSO-128) were grown in M9 Glu^+^ to OD_600_ ~0.5 and two 0.5 mL subcultures were prepared for each culture. The use of D23580 ΔΦ [P22] in these experiments controlled for the effect of lysis from without due to use of high multiplicity of infection (MOI). The strain is a lysogen for WT P22 phage, and therefore is highly resistant to infection by P22-derived phages. P22 Δ*c2* was added at an MOI of 5. The same volume of LB was added to the two remaining subcultures (non-infected controls). Samples were incubated for 15 min at 37°C to allow phage attachment. To stop phage development, the cultures were chilled on ice and bacteria were washed with 0.5 mL of cold PBS. All the samples were serial-diluted in PBS to 10^-6^ and kept on ice. For the measure of survival post-infection, 10 μL of diluted infected or non-infected cultures were applied in technical triplicates on LB agar supplemented with 10 mM EGTA (EGTA was used to minimize secondary infection by free phages). Colony forming Units (CFU) were enumerated and the survival rate, was calculated as the ratio of CFUs in infected cultures divided by the CFUs obtained from non-infected cultures (in %). All the survival experiments were carried out at least twice with biological triplicates.

### Frequency of lysogeny assays

For the frequency of lysogeny assays, a derivate of phage P22 was used that has the *pid* locus replaced with an *aph* cassette yielding kanamycin resistant lysogens (P22 *Δpid::aph*, SNW490). The *pid* locus has previously been shown to be non-essential in phage P22 and does not establishment of lysogeny (Cenens et al., 2013). D23580 ΔΦ *tetR-P_tetA_-bstA* (SNW576) cells were grown in 3 mL of LB to OD_600_ ~0.35. Methanol (mock) or AHT (500 ng/mL, inducer) were added to the cultures and bacteria were incubated to induce BstA for 1 hour at 37°C. 200 μL samples of the bacteria were mixed in triplicate with P22 *Δpid::aph* phage to achieve a MOI of 0.1, and incubated at 37°C for 20 minutes to allow adsorption and ejection of nucleic acids. Cells were pelleted and resuspended in LB media supplemented with 10 mM EGTA to minimize secondary infection by any free phages (along with methanol or AHT) and incubated at 37°C for a further 20 minutes to allow integration and expression of the kanamycin resistance determinant. Colony forming Units (CFU) were enumerated on LB kanamycin. Frequency of lysogeny was determined as the kanamycin resistant CFU/mL divided by the PFU/mL of input phage.

### Phage DNA detection by Southern Blotting

D23580 ΔΦ *tetR-P_tetA_-bstA* (SNW576) was grown in 50 mL LB to OD_600_ ~0.35. The culture was split in two 20 mL sub-cultures and methanol (mock) or AHT (500 ng/mL, inducer) were added to each subculture. Bacteria were incubated to induce BstA for 20 min at 37°C and the phage of interest was added at an MOI of 5. Infections were carried out at 37°C with aeration and total DNA was extracted (Quick-DNA™ Universal Kit Zymo) from 1.5 mL of culture at 0, 10, 20, 30, 35, 40 and 50 minutes post Infection. Total DNA (100 ng, according to QuBit quantification) was size-separated (2 hours at 100 V in TAE 1X) on a 0.8 % agarose-TAE gel containing Midori Green DNA staining (4 μL for 100 mL gel). One hundred nanograms of none-infected D23580 ΔΦ *tetR-P_tetA_-bstA* genomic DNA were used as a negative control. DNA was fragmented by exposing the agarose gel to UV light for 5 min on a UV-transilluminator. DNA was denatured by soaking the gel in the Denaturation Solution (0.5 M NaOH, 1.5 M NaCl) for 30 min and then in the Neutralization Solution (1.5 M NaCl, 1 M Tris-HCl, pH 7.6) for 30 min. DNA was transferred on a positively-charged Nylon membrane using the capillary blotting method. Phage DNA was detected with DIG labelled dsDNA probes generated by PCR amplification with MyTaq DNA polymerase (Bioline), buffer, phage DNA and primers (0.4 μM each), in the presence of 0.2 mM dATP, 0.2 mM dCTP, 0.2 mM dGTP, 0.13 mM dTTP and 0.07 mM DIG-11-dUTP. For the 9NA probe a 588 bp PCR fragment was generated with primer pair NW_602 / NW_603 and for the P22/BTP1 probe a 725 bp PCR fragment was generated with primer pair SO-22 / SO-23. The DNA probes were heat-denatured at 95°C for 15 min and the DNA-DNA hybridizations were carried out at 45°C for 16 hours in DIG-Easy Hyb buffer. The washing and immunodetection procedures were carried out, as specified in the DIG Application Manual for Filter Hybridization (Roche) and the chemilumiscence signal was detected using an ImageQuant LAS 4000 imager (GE Healthcare). Prior to DNA transfer onto the membrane, the Midori green-stained DNA was visualized under UV and the resulting image was used as a loading control.

### Phagemid efficiency of transformation

To avoid a reduction in transformation caused by interference interspecies DNA modification/restriction interference between *E. coli* and *Salmonella*, all the P22-derived phagemids were replicated and extracted from *S.* Typhimurium SNW555.

*Salmonella* strains carrying the *tetR-P_tetA_-bstA* module were grown in 50 mL LBO culture. When OD_600_ ~0.4 was reached, each culture was split into two 25 mL sub-cultures and methanol (mock) or AHT (inducer) were added to each subculture. Bacteria were incubated for BstA induction during 15 min at 37°C. The cultures were incubated on ice for 5 min and bacteria were washed twice with cold water (25 mL) and were concentrated in 0.1 mL of ice-cold sterile 10% glycerol. The OD_600_ of each electro-competent cell sample was measured by diluting 10 μL of competent cells with 990 μL of 10% glycerol. Cell concentration was adjusted with 10% glycerol for each sample, according to the sample with the lowest OD_600_. The competent cells (20 μL) were mixed with 10 ng (estimated by Qubit) of the P22 phagemids, pP22 (pNAW229) or pP22-*aba* (pNAW230) and the mixture was incubated on ice until electroporation (2.5 KV). Transformation reactions were re-suspended in 1 mL LB or 1 mL LB + AHT (for the *bstA*-induced bacteria) and were incubated for 60 min at 37°C, for recovery. The transformations were diluted (decimal dilution to 10^-5^) in LB or LB+AHT and 100 μL of each dilution (including the non-diluted sample) were spread on LB agar Km or LB agar Km+AHT plates. After incubation at 37°C, the number of Km^R^ transformants was enumerated for each transformation and efficiency of transformation was defined as the number of transformants obtained per ng of phagemid. This experiment was performed with biological triplicates and was repeated twice with LT2 *tetR-P_tetA_-bstA* (SNW389) and once with D23580 ΔΦ *tetR-P_tetA_-bstA* (SNW576), giving similar results.

### Microscopy-general

For all imaging experiments, bacteria were sub-cultured in liquid M9 Glu^+^ media. All images were collected with a wide field Nikon Eclipse Ti-E inverted microscope equipped with an Okolab Cage Incubator warmed to 37°C with Cargille Type 37 immersion oil. A Nikon CFI Plan Apo DM Lambda 100X 1.45 NA Oil objective and a Nikon CFI Plan Apo DM Lambda 20X .75 NA objective were used with Perfect Focus System for maintenance of focus over time. Superfolder GFP, mCherry and SYTOX Orange Nucleic Acid Stain (ThermoFisher) were excited with a Lumencor Spectra X light engine with Chroma FITC (470/24) and mCherry (575/25) filter sets, respectively and collected with a Spectra Sedat Quad filter cube ET435/26M-25 ET515/30M-25 ET595/40M-25 ET705/72M-25 and a Spectra CFP/YFP/mCherry filter cube ET475/20M-25 ET540/21M-25 ET632/60M-25. Images were acquired with an Andor Zyla 4.2 sCMOS controlled with NIS Elements software. For time-lapse experiments, images were collected every 3 minutes (unless specified otherwise) *via* ND acquisition using an exposure time of 100 ms and 50% or 100% illumination power for fluorescence. Multiple stage positions (fields) were collected using the default engine Ti Z. Fields best representing the overall experimental trend with the least technical artefacts were chosen for publication. Gamma, brightness, and contrast were adjusted (identically for compared image sets) using FIJI(Schindelin et al., 2012). The FIJI plugins Stack Contrast (Capek et al., 2006) and StackReg (Thevenaz et al., 1998) were used for brightness matching and registering image stacks.

### Microscopy-agarose pads

Agarose pads were prepared with 2% agarose and M9 Glu^+^ media, and mounted on MatTek dishes (No. 1.5 coverslip, 50 mm, 30 mm glass diameter, uncoated). Cells (D23580Δ*tsp-gtrAC* (JH4287) or *D23580Δtsp-gtrAC bstA^STOP^* (SNW431) were grown to log phase (OD_600_ ~ 0.4) in M9 Glu^+^ at 37°C with shaking (220 RPM), and where required, diluted in fresh M9 Glu^+^ to achieve the desired cell density on the agarose pad. For experiments where all cells were infected (Figure 4A), phage P22 Δ*c2* was added at an MOI of 5. Phage adsorption and initial infection was facilitated by incubation at 37°C with shaking for 10 minutes. Subsequently, infected cells were pelleted at 5000 x g and resuspended in ice-cold PBS to pause phage development. Two microliters of chilled, infected cells were spotted onto opposite sides of an agarose pad (two strains were imaged on the same pad) and inverted onto the MatTek imaging dish. Experimental MOIs were immediately confirmed by CFU and PFU /mL measurement of the cell and phage preparations. Phase-contrast images using the 100X objective were collected every 3 minutes for 3 hours.

Procedures for experiments involving a subset of infected cells (Figure 4C) were identical, except cells infected with P22 Δ*c2 P-mcherry* were washed an additional 4 times in ice-cold PBS to reduce the concentration of un-adsorbed, free phage. In parallel, uninfected cells were washed once in ice-cold PBS. Infected cells were mixed at a ratio of 1:1000 with uninfected cells of the same genotype before being spotted onto the agarose pad. For these experiments, phase-contrast and fluorescence images (mCherry) using the 20X objective were collected every 3 minutes for 3 hours.

### Microscopy-microfluidic infection

The CellASIC ONIX2 system from EMD Millipore with B04A plates was used for microfluidic imaging experiments (Figure 5). Phages used in microfluidic infection experiments shown in 5B (P22 HT or 9NA) were stained with SYTOX Orange Nucleic Acid Stain according to the protocol previously described (Valen et al., 2012). Stained phages washed 4 times in 15 mL M9 Glu^+^ media using Amicon Ultra-15 centrifugal filter units. After staining, the titer and viability of phages were immediately assessed by plaque assay, and once stained, phages were used for no longer than 2 weeks. For use in the microfluidic experiments, SYTOX Orange strained phages were normalised to a titer of approximately 10^10^ PFU/mL. Cells (D23580 *bstA-_sf_gfp*, SNW403) were grown to early exponential phase (OD_600_ ~ 0.1) in M9 Glu^+^ at 37°C with shaking (220 RPM) before being loaded into CellASIC B04A plates using the pressure-driven method according to the manufacturer protocol for bacterial cells. The slanted chamber of the plate immobilises the cells, but allows media to flow continuously. Firstly, cells were equilibrated with constant M9 Glu^+^ media flow for approximately 30 minutes. Secondly, stained phages suspended in M9 Glu^+^ media were flowed over the cells until the majority of cells were infected (typically 10-30 minutes). In the case of P22 HT phage (which exhibits inefficient adsorption to D23580 *bstA-_sf_gfp* due to the *gtr* locus of prophage BTP), phages were continuously flowed. Finally, M9 Glu^+^ media was flowed over the cells for the duration of the experiment. Microfluidic experiments typically lasted 5 hours, after which time uninfected cells outgrew the chamber. Phase-contrast and fluorescence images were collected every 1.5 minutes for the experiments in Figure 5B.

For the microfluidic imaging experiments shown in Figure 5C, strain SVO251 (S. Typhimurium D23580 ΔΦ *STM1553::(P_tetA_-bstA-_sf_gfp-frt*) ΔpSLT-BT ΔpBT1 pAW61 (*P_BAD_-parB-mcherry*) was used. This strain contains the *bstA-_sf_gfp* fusion contrast under the control of the *P_tetA_* promoter. However, this strain lacks the *tetR* gene, and therefore expression of *bstA-_sf_gfp* is constitutive (not inducible). Additionally, this strain is cured of two natural plasmids that contain native partitioning systems (pSLT-BT and pBT1), and there for might interfere with the correct function of the ParB-*parS* system used for phage DNA localization. The ParB-mCherry fusion protein is expressed from the pAW61 plasmid (Ap^R^) under the control of the P_BAD_ promoter (induced by L-arabinose). Strain SVO251 was grown in M9 Glu^+^ supplemented with 100 μg/mL ampicillin to maintain the pAW61 plasmid and 0.2% L-arabinose to induce expression of ParB-mCherry. The same supplemented media was used in the microfluidic chamber. Cells were grown to ~OD_600_ 0.1 before loading into the CellASIC B04A plate as described above. After 15 minutes growth, phage P22 *Δpid::(parS-aph)* [which contains one parS site along with a kanamycin resistance locus, *aph*, in place of the non-essential *pid* locus (Cenens et al., 2013)] diluted to a concentration of 10^8^ PFU/mL (in M9 Glu^+^ amp100 0.2% L-ara) was flowed into the chamber. Phase contrast and red and green fluorescence images were collected every 2 minutes for 4 hours.

### BstA protein homolog analysis

BstA protein homologs were identified using tblastn (database: non-redundant nucleotide collection) and the HMMER webserver (Potter et al., 2018) (database: Reference Proteomes). The dataset of BstA protein homologs was manually curated to reflect the diversity of taxonomic background harbouring homologs. Evolutionary covariance analysis was done using DeepMetaPSICOV (Buchan and Jones, 2019) at the PSI-PRED server (Kandathil et al., 2019). To analyse the genetic context of BstA homologs, the sequence region 20 Kb either side of the homolog (40 kb total) was extracted (BstA 40 kb neighbourhoods). To produce homogenous and comparable annotations, each region was re-annotated using Prokka 1.13 (Seemann, 2014). Additionally, the resulting annotated amino acid sequences were queried against our custom BstA profile-hmm and the Pfam 31.0 database (El-Gebali et al., 2019) with hmmerscan (Eddy, 1998), and the highest scoring significant hit per ORF was considered for the results shown in Figure 2. All the code is available in https://github.com/baymLab/2020_Owen-BstA.

Pairwise identity of homologs in Figure 2B to BstA^BTP1^ was computed using the EMBOSS Needle webserver (Needleman and Wunsch, 1970). BstA homologs were designated “putatively-prophage associated” if annotated genes in the 40 kb neighbourhood contained any instance or the word “phage” or “terminase”. For categorisation in Figure 2C, homologs were classed as having “high confidence association” if instances of gene annotations including the aforementioned key words occurred both before, and after, the BstA gene within the 40 Kb neighbourhood (i.e., to account for the possibility that a prophage-independent homolog could co-occur next to a prophage region by chance). Homologs classed as having “low confidence association” had at least one instance of genes whose annotations included “phage” or “terminase” either in the upstream or downstream 20Kb, but not both. Plasmid status was determined from information in the sequence records. The HHpred webserver was used to annotate the putative KilA-N domain (Zimmermann et al., 2018). All homolog neighbourhoods, homolog alignments and sequences is available to download https://github.com/baymLab/2020_Owen-BstA.

## QUANTIFICATION AND STATISTICAL ANALYSES

The phage replication, survival rate, efficiencies of transformation and of lysogeny were calculated as mentioned above. The numerical data were plotted and analyzed using GraphPad Prism 8.4.1. Unless stated otherwise in the figure legends, data are presented as the mean of biological triplicates ± standard deviation. The unpaired *t*-test was used to compare the groups and statistical significance is indicated on the figures. P values are reported using the following criteria: < 0.0001 = ****, 0.0001 to 0.001 = ***, 0.001 to 0.01 = **, 0.01 to 0.05 = *, ≥ 0.05 = ns.

### KEY RESOURCES TABLE

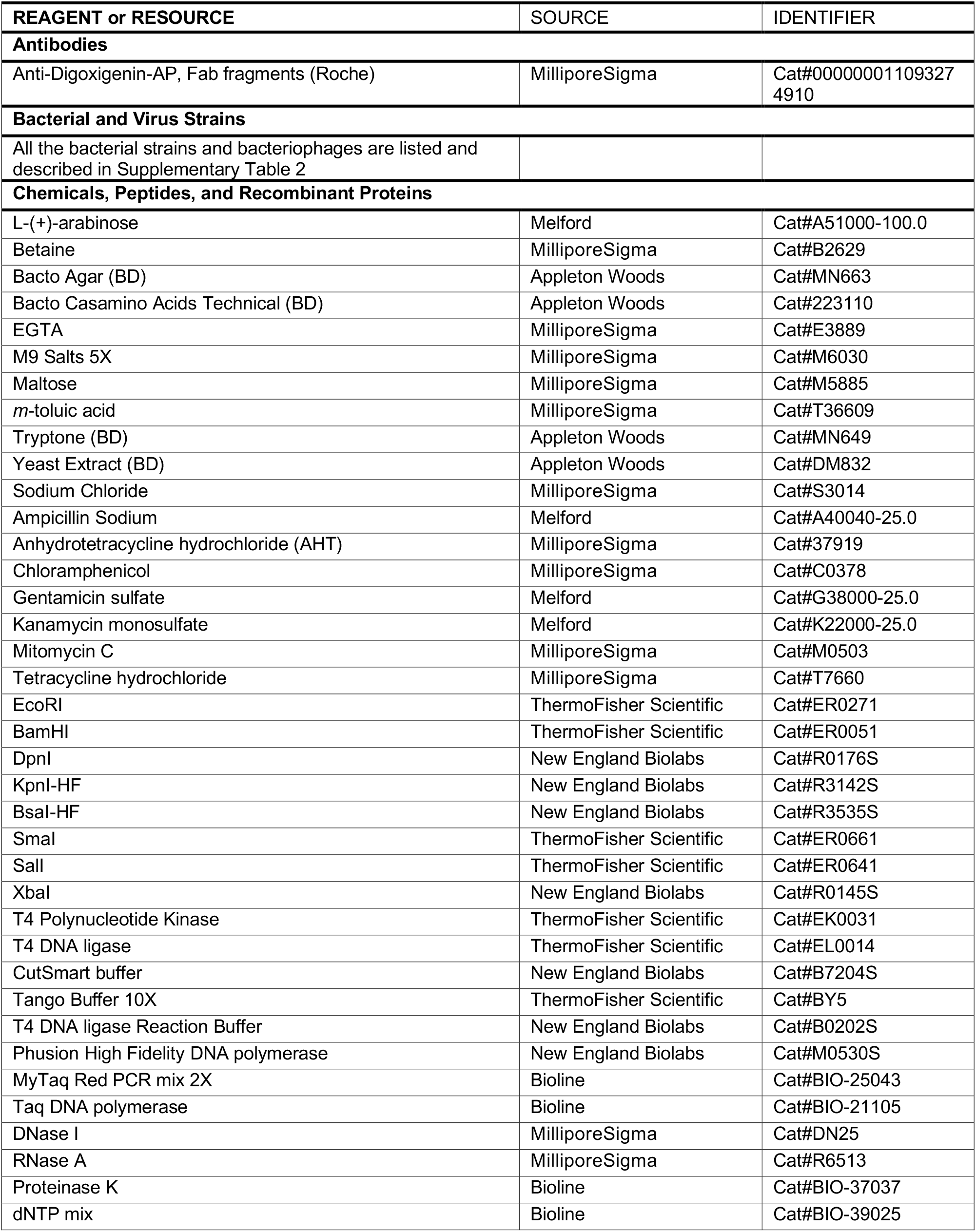
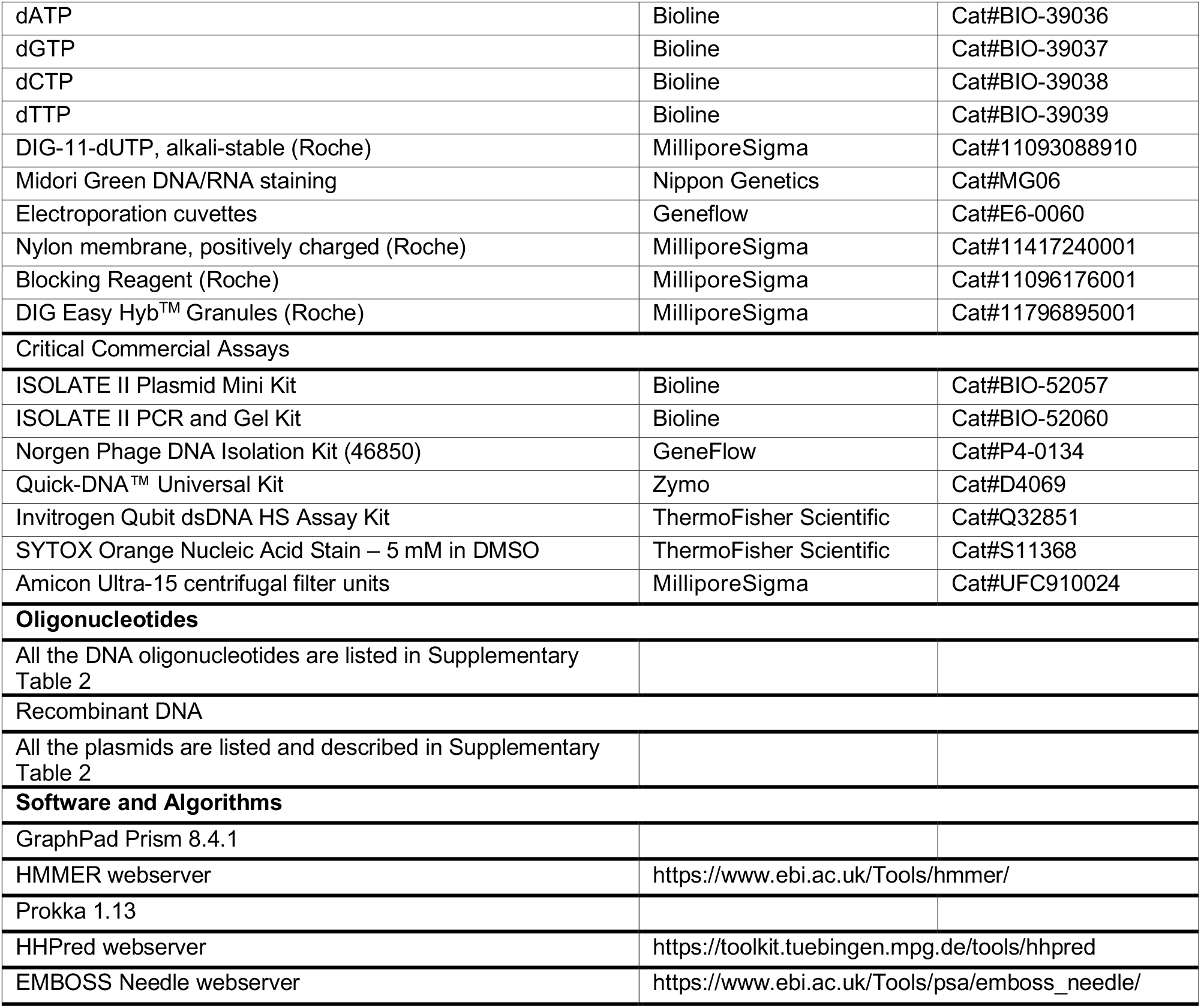

## Acknowledgements

We are grateful to present and former members of the Hinton and Baym laboratories for helpful discussions, and Paul Loughnane for his expert technical assistance. We thank the Bollback Lab (Liverpool) for the gift of pCas9 and P1*vir*, Allison Lab (Liverpool) for MG1655 and T4, Casadesus Lab (Seville) for 9NA and Det7, Ansaldi Lab (Marseille) for T5, Penadés Lab (Glasgow) for ES18, and Klumpp Lab (Zurich) for FelixO1. We also acknowledge all the other generous researchers who responded to our call for *Salmonella* and *E. coli* phages. We thank Benoît Doublet for the gift of pCP20-Gm, and the Van Valen Lab, Francois St-Pierre and Paul Wiggins for λFS135 and pAW62. We thank Thomas K. Wood for helpful comments.

Research in this publication was supported by a Wellcome Trust Senior Investigator award (to JCDH, grant number 106914/Z/15/Z), NIGMS of the National Institutes of Health (to MB, award number R35GM133700) and the David and Lucile Packard Foundation (SVO, NQO, and MB).

## Figure legends & Supplementary Materials

**Supplementary Table 1**: Details of all BstA homologs used in the analysis presented in Figure 2.

**Supplementary Table 2**: Details of all oligonucleotide sequences, plasmids, bacterial strains and phages used in this study.

**Supplementary Video 1**: Cells natively expressing BstA (D23580 *Δtsp-gtrAC,* JH4287) or possessing a mutated BstA locus (D23580 *Δtsp-gtrAC bstA^STOP^*, SNW431) were infected with the obligately virulent P22-derivate phage, P22 Δ*c2*, at an MOI of 5 (to increase the likelihood of infecting all cells). Infected cells were imaged every 3 minutes on agarose pads. Regardless of BstA function, almost all cells were observed to lyse (indicated by loss of defined cell shape and phase contrast).

**Supplementary Video 2**: Cells natively expressing BstA (D23580 Δ*tsp-gtrAC*, JH4287) or possessing a mutated BstA locus (D23580 *Δtsp-gtrAC bstA^STOP^*, SNW431) were infected with the obligately virulent P22-derivate phage, P22 Δ*c2 P-mcherry*, at an MOI of 5 (to increase the likelihood of infecting all cells). Due to the *mcherry* insertion, red fluorescence corresponds to phage replication. Infected cells were mixed 1:1000 with uninfected cells. Cells mixtures were imaged every 5 minutes on agarose pads for 6 hours. In the BstA+ cells, primary infected cells lysed, but did not stimulate secondary infections of neighbouring cells, and eventually formed a confluent lawn. In BstA-cells, primary lysis events caused secondary infections (neighbouring cells showing red fluorescence and subsequent lysis) causing an epidemic of phage infection reminiscent of plaque formation.

**Supplementary Video 3**: Cells natively expressing BstA translationally fused to _sf_GFP (D23580 *bstA-_sf_gfp*, SNW403) were grown in a microfluidic growth chamber and imaged every 1.5 minutes. Fluorescently labelled phages P22 HT (left) or 9NA (right) were then added to the cells, and can be seen adsorping to cells as red fluorescent puncta. For purposes of comparison, timestamps are synchronised to the point at which phage are first observed. Typically around 20 minutes after initial observation of phage infection, BstA proteins formed discrete and dynamic foci within the. Cells then proceeded to lyse.

**Supplementary Video 4**: Cells heterologously expressing BstA translationally fused to _sf_GFP, and ParB translationally fused to mCherry (D23580 ΔΦ ΔpSLT-BT ΔpBT1 *STMMW_15481::[P_tetA_-bstA^BTP1^-_sf_gfp-frt]* pAW61, SVO251) were grown in a microfluidic growth chamber and imaged every 2 minutes. Phage P22 *Δpid::(parS-aph)* were then added to the cells. ParB protein oligomerises onto DNA at *parS* sites, and therefore *parS*-tagged DNA is indicated by ParB-mCherry foci (red). Translocation of infecting phage DNA into cells is indicated by the formation of red foci within the cells, which is rapidly followed by formation of green, BstA foci. The green and red foci appear to physically overlap. Infected cells proceed to lyse. The same timeseries is presented as separate channels (phase contrast, GFP, mCherry), and a composite merge.

---

**Supplementary Figure 1:**
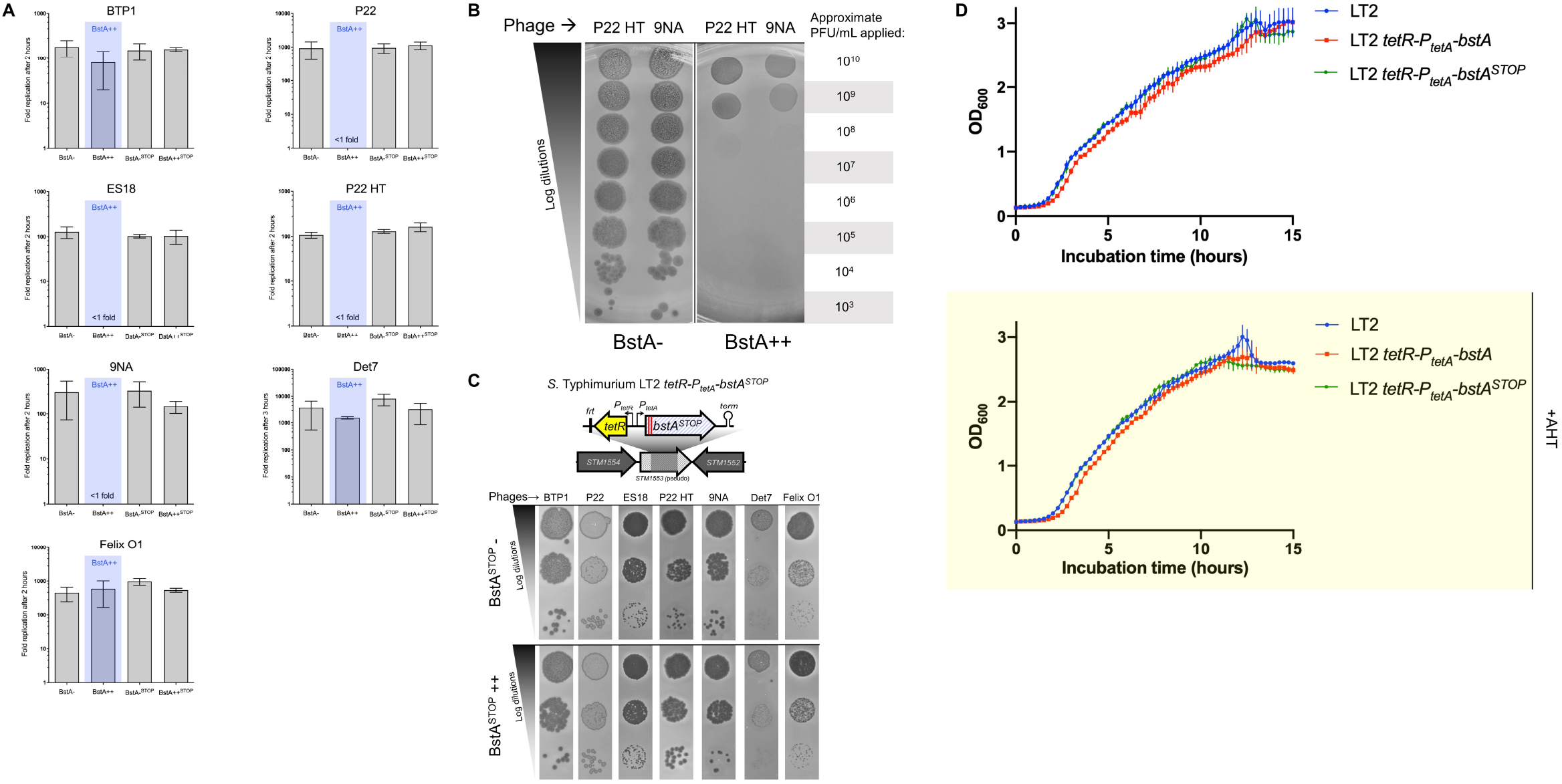
The *bstA* locus confers phage resistance and *bstA* nonsense mutations supresses the phage resistance phenotype. (**A**) Replication assays of the indicated phages were carried out with mock-induced (BstA-) or AHT-induced (BstA++) strains LT2 *tetR-P_tetA_-bstA^BTP1^* (JH4400) or LT2 *tetR-P_tetA_-bstA^STOP^* (JH4402, carrying two nonsense mutations in *bstA*), as host. Phage replication was measured 2-3 hours post infection and phages were enumerated on lawns of LT2 WT. Phage replication is presented as the mean of biological triplicates ± SD. (**B**) Extended concentration plaque assay of phage P22 HT and 9NA on LT2 *tetR-P_tetA_-bstA^BTP1^* (JH4400) with mock (BstA-) or AHT induction (BstA++). The approximate PFU/mL (**C**) Nonsense mutations in *bstA* suppress the BstA-driven anti-phage phenotype. Plaque assays were carried out with the indicated phages on mock-or AHT-induced lawns of LT2 *tetR-P_tetA_-bstA^STOP^* (JH4402). (**D**) Optical density growth curves (OD_600_) of LT2 WT, LT2 *tetR-P_tetA_-bstA^BTP1^* (JH4400) or LT2 *tetR-P_tetA_-bstA^STOP^* (JH4402, carrying two nonsense mutations in *bstA*)) with and without AHT induction. No cellular toxicity (as measured by reduced culture growth) was associated with *bstA^BTP1^* or *bstA^STOP^* expression.

**Supplementary Figure 2:**
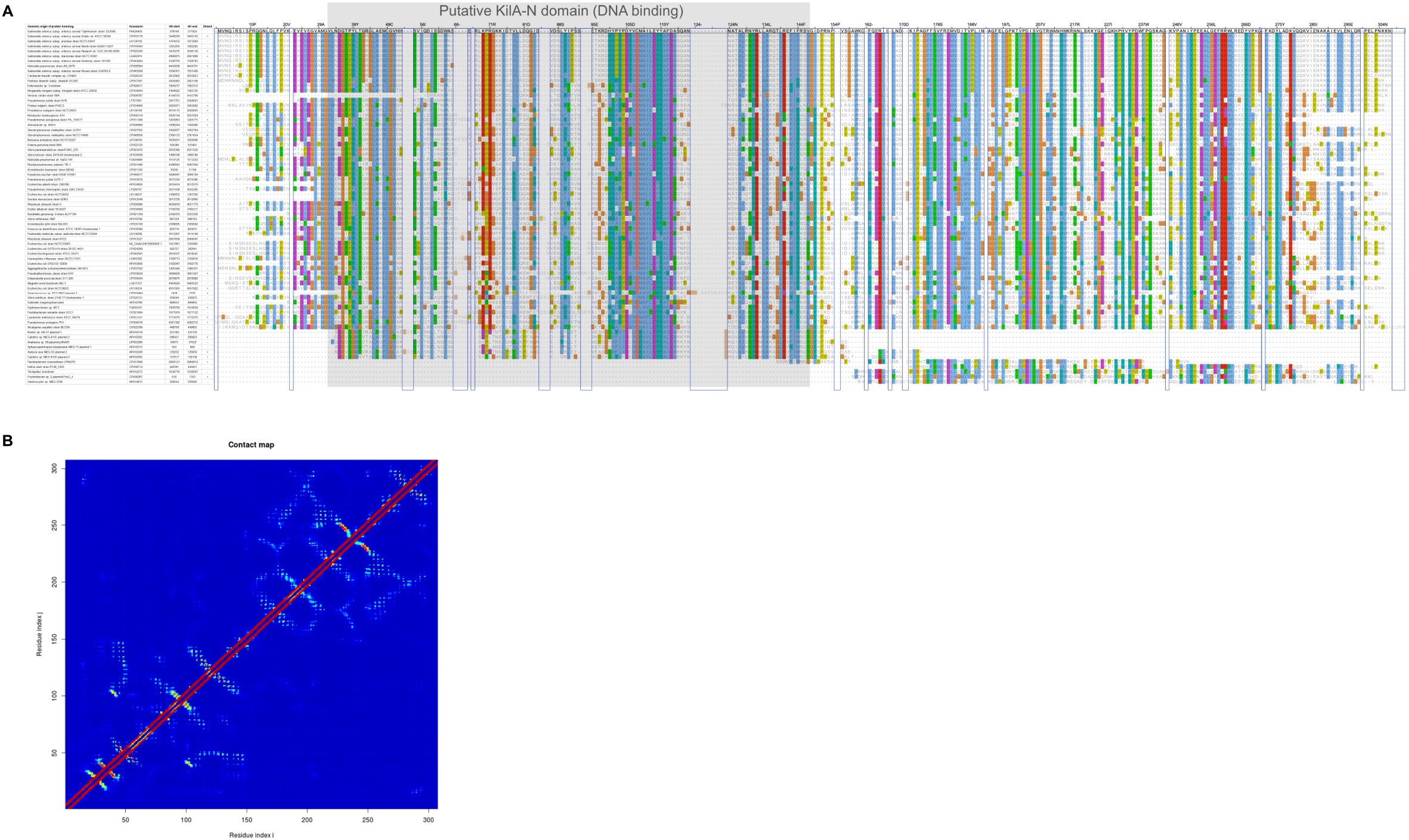
Extended alignment of BstA protein homologs and predicted BstA contact map. (A) Blue boxes indicate columns which are gaps relative to the reference sequence (top row, BstA from *S.* Typhimurium D23580), and are collapsed in the alignment shown in Figure 2. Grey box indicates the position of the putative KilA-N domain. (B) Predicted contact map derived by evolutionary covariance analysis of BstA by DeepMetaPSICOV. The confidence of a pair of residues x and y interacting is indicated by the colour of the cell at position x, y (and by mirroring at position y,x) with red indicating high probability of interaction and dark blue indicating low probability. The map suggests that residues 1-~155 and ~156-end form folded domains that interact very little.

**Supplementary Figure 3:**
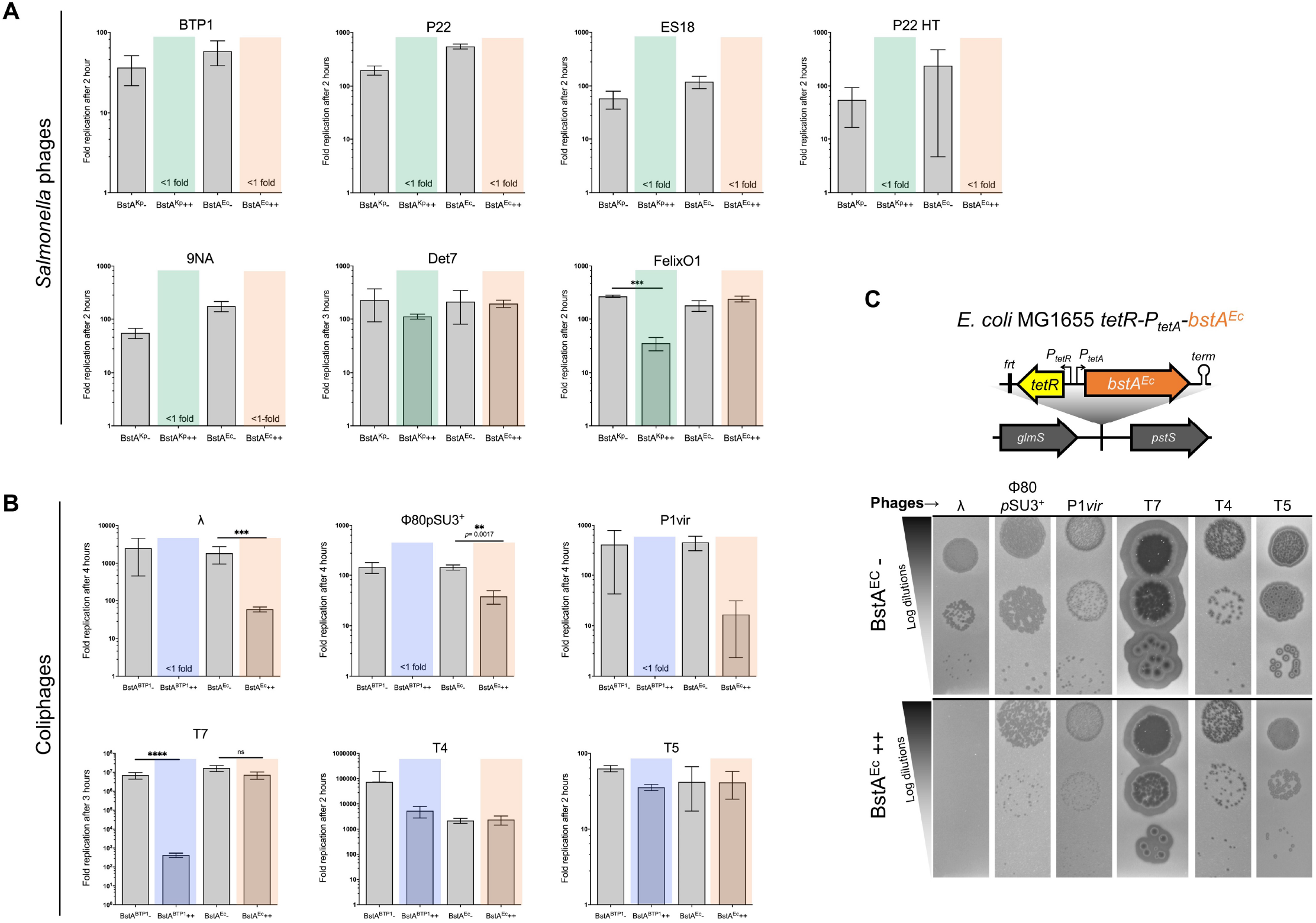
BstA homologs confer phage resistance in *S.* Typhimurium LT2 and *E. coli* MG1655. (**A**) BstA^Kp^ and BstA^Ec^ confer phage resistance in *S.* Typhimurium. (**B**) BstA^BTP1^ and BstA^Ec^ confer phage resistance in *E. coli*. Phage replication assays were carried out with mock-induced (BstA-) or AHT-induced (BstA++) cultures of LT2 *tetR-P_tetA_-bstA^Ec^* (JH4408), LT2 *tetR-P_tetA_-bstA^Kp^* (JH4404), MG1655 *tetR-P_tetA_-bstA^BTP1^* (JH4410) or MG1655 *tetR-P_tetA_-bstA^Ec^* (JH4414), infected with the indicated phage. LT2 or MG1655 WT lawns were used for phage enumerate 2-3 hours post infection. Phage replication is presented as the mean of biological triplicates ± SD. When replication difference between induced and noninduced cultures was lower than one order of magnitude, groups were compared using unpaired two-tailed Student *t*-test and P values and significance are indicated by *,**,*** or ns (not significant). (**C**) BstA^Ec^ confers phage resistance to *E. coli*. Plaque assay were carried out with the indicated phages applied on a lawn of mock-or AHT-induced MG1655 *tetR-P_tetA_-bstA^Ec^* (JH4414).

**Supplementary Figure 4:**
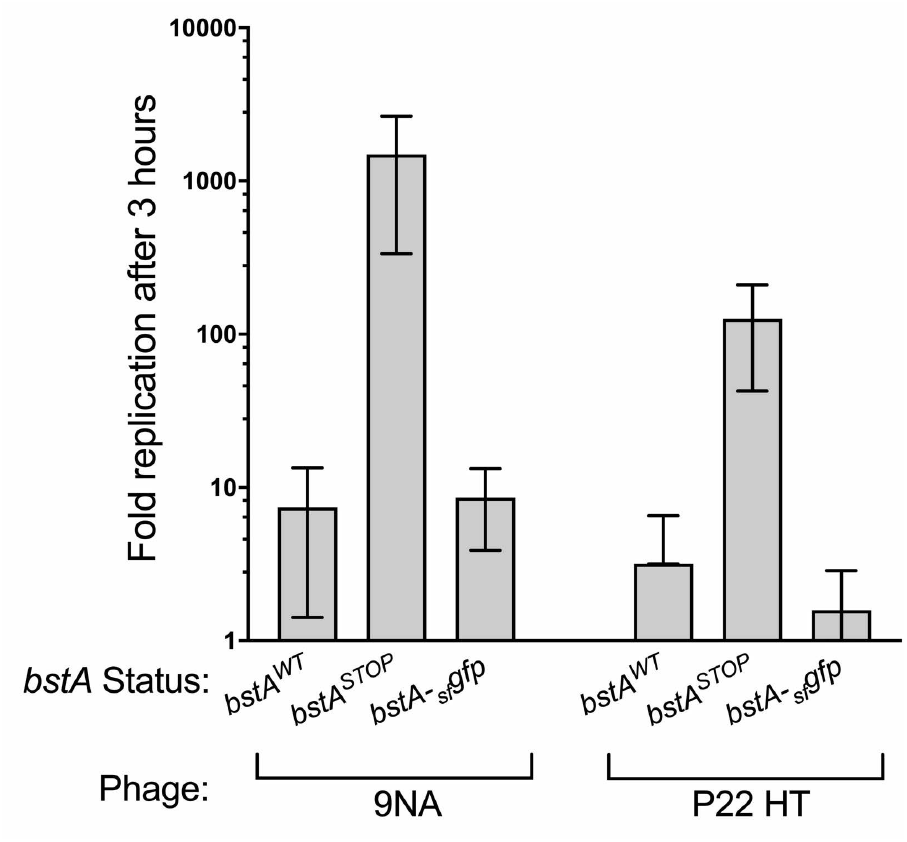
C-terminal Superfolder GFP (_sf_GFP) fusion to BstA does not impair the BstA anti-phage activity. Replication assays were carried with the indicated phages on strains carrying the WT *bstA (bstA^WT^*, strain D23580 in panel A, the defective *bstA^STOP^* version (strain SSO-78) or the BstA-sfGFP fusion strain (*bstA-_sf_gfp*, strain SNW403). Phage replication was measured 3 hours post-infection and plaques were enumerated on lawns of D23580 *Δtsp-gtrAC bstA^STOP^* (SNW431). Phage replication is presented as the mean of biological triplicates ± SD.

**Supplementary Figure 5:**
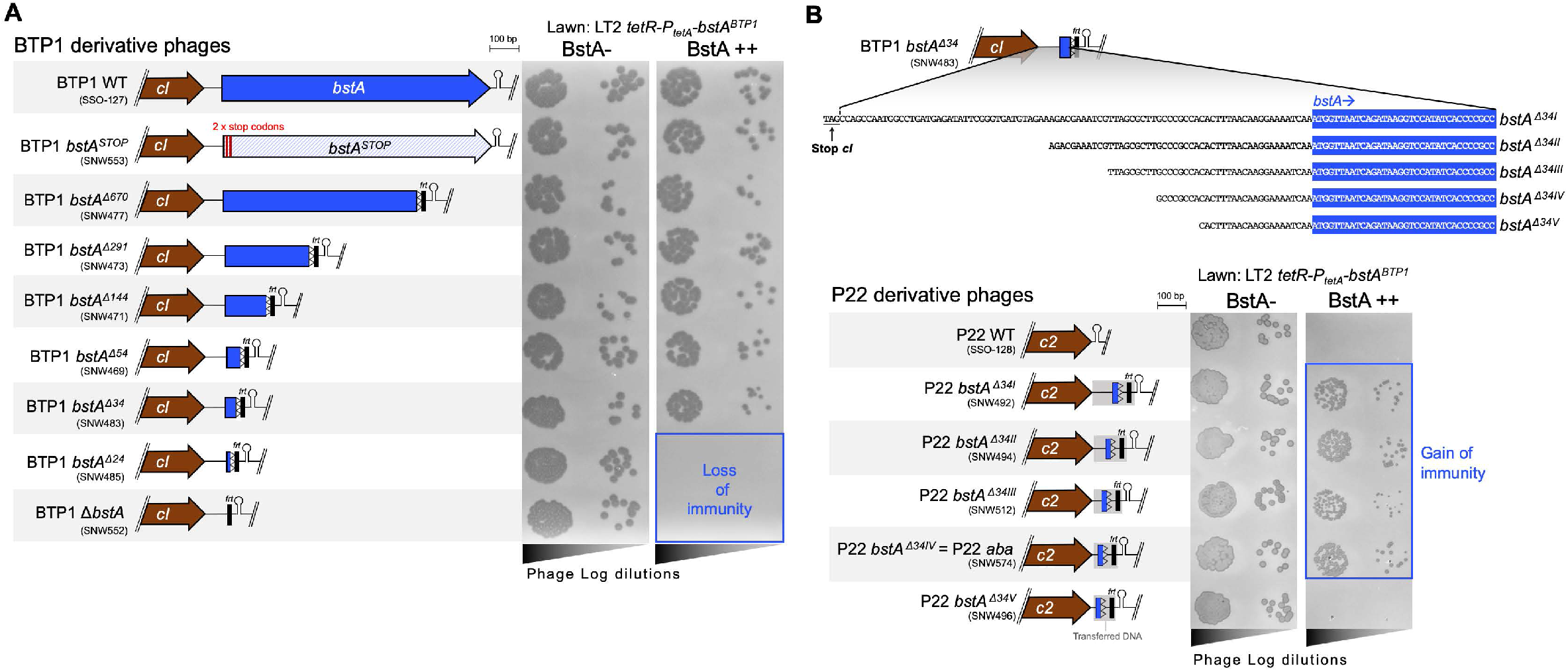
Identification of the anti-BstA (*aba*) factor. (**A**) BTP1 *bstA* locus truncations revealed the location of the *anti-bstA* fragment *aba*. The *bstA* truncations are indicated for each BTP1 variant. The *bstA^Δ^* allelic numbering corresponds to the number of base pairs remaining from the *bstA* ATG start. (**B**) Transfer of the *bstA^Δ34^* fragment in P22 confers BstA-immunity. Fragments *bstA^Δ34I^-bstA^Δ34V^* are indicated. The donor lysogen strain for each BTP1 and P22 variant is indicated in brackets. The residual scar sequence of pKD4 is indicated by *frt* and hairpins represent Rho-independent terminators. Plaque assays were performed with the indicated BTP1/P22-derived phages on lawns of mock-induced (BstA-) or AHT-induced (BstA++) LT2 *tetR-P_tetA_-bstA^BTP1^* (JH4400).

**Supplementary Figure 6:**
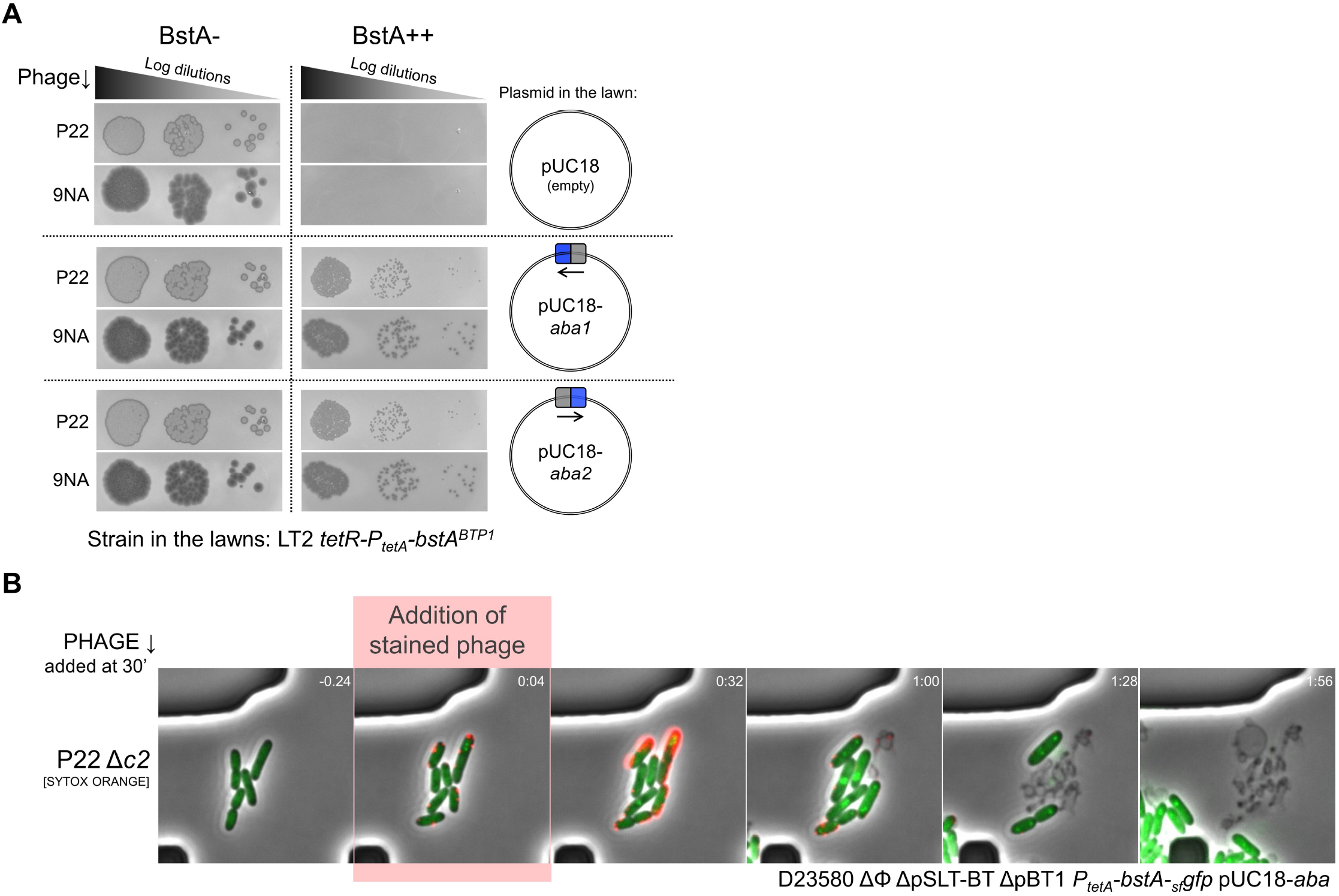
Multiple copies of *aba* DNA suppresses BstA activity in *trans* but do not affect protein localization dynamics. (**A**) The *aba* sequence was cloned into the pUC18 plasmid in either orientation, and the plasmids were transformed into LT2 *tetR-P_tetA_-bstA^BTP1^* (JH4400). Plaques assays were performed with phage P22 and 9NA applied on lawns of mock-induced (BstA-) or AHT-induced (BstA++) LT2 *tetR-P_tetA_-bstA^BTP1^* (JH4400), transformed with the indicated plasmid. Both plasmids (pUC18-*aba1* and pUC18-*aba2*) supressed BstA activity and permitted plaque formation by P22 and 9NA in the presence of BstA. (**B**) A microfluidic growth chamber was used to observe the behaviour of BstA protein during phage infection in the presence of the pUC18-*aba* plasmid, capturing images every 4 minutes. A time series of representative fields are presented as composite images (phase contrast, green and red fluorescence are overlaid). Cells (D23580 ΔΦ ΔpSLT-BT ΔpBT1 *P_tetA_-bstA-_sf_gfp* pUC18-*aba*, SVO254) were first grown for a period in the chamber (immobilised by the angle of the chamber ceiling) with constant flow of M9 Glu^+^ amp100 media (Methods). Fluorescently labelled phage P22 Δ*c2* (stained with SYTOX Orange resuspended in M9 Glu^+^ media, Methods) were then added to the cells. For purposes of comparison, timestamps are synchronised to the point at which phage are first observed adsorbing to cells. Localisation of BstA proteins into foci proceeding cell lysis was conserved in the presence of the pUC18-*aba* plasmid.

**Supplementary Figure 7.**
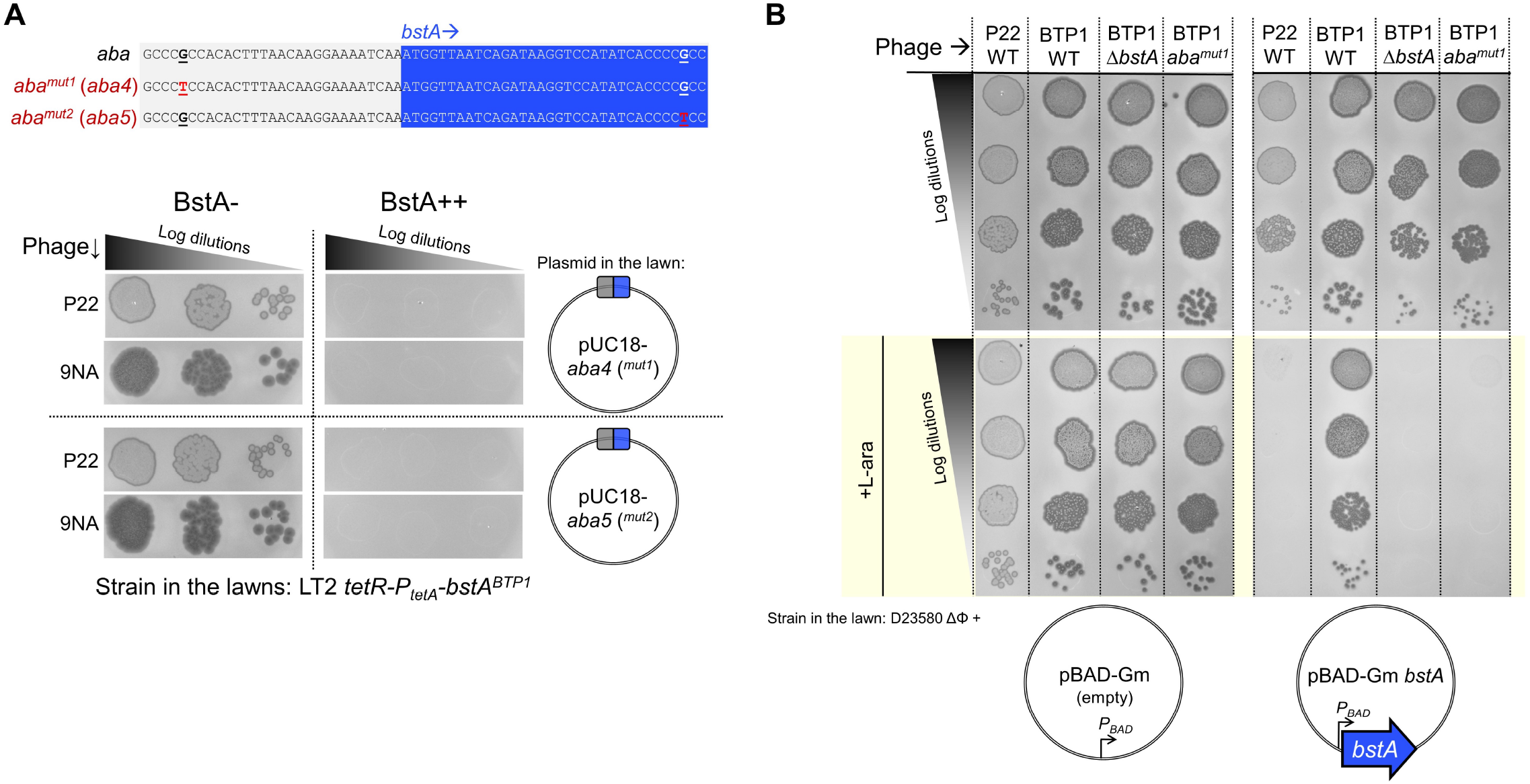
Mutations in the terminal direct repeat motifs of *aba^BTP1^* ablate its function in *trans* and during phage infection. (**A**) The assay shown in Supplementary Figure 6A was repeated but with mutated versions of the *aba* sequence. When a single nucleotide mutation was made in either of the terminal CCCGCC motifs, the pUC18 *aba* plasmids could not rescue the replication of P22 or 9NA in the presence of BstA. (**B**) BTP1 phage carrying the *aba^mut1^* mutation (which does not alter the coding sequence of the *bstA* gene), were challenged against cells expressing BstA from an arabinose-inducible promoter (*P_BAD_*) on a plasmid (plasmid pNAW254). Phages BTP1 WT, P22 WT and BTP1 *ΔbstA* (in which *aba*-mediated BstA immunity is present, absent, and synthetically absent, respectively) were included as comparators. BTP1 *aba^mut1^* plaques normally in the absence of BstA (in cells carrying the empty plasmid, or cells harbouring the *bstA* plasmid without +L-ara inducer). BTP1 *aba^mut1^* is unable to form plaques on cells expressing BstA, and is inhibited to the same degree as phage P22 (entirely lacking *aba*), and BTP1 *ΔbstA* (in which half of the *aba* sequence is deleted).

**Supplementary Figure 8.**
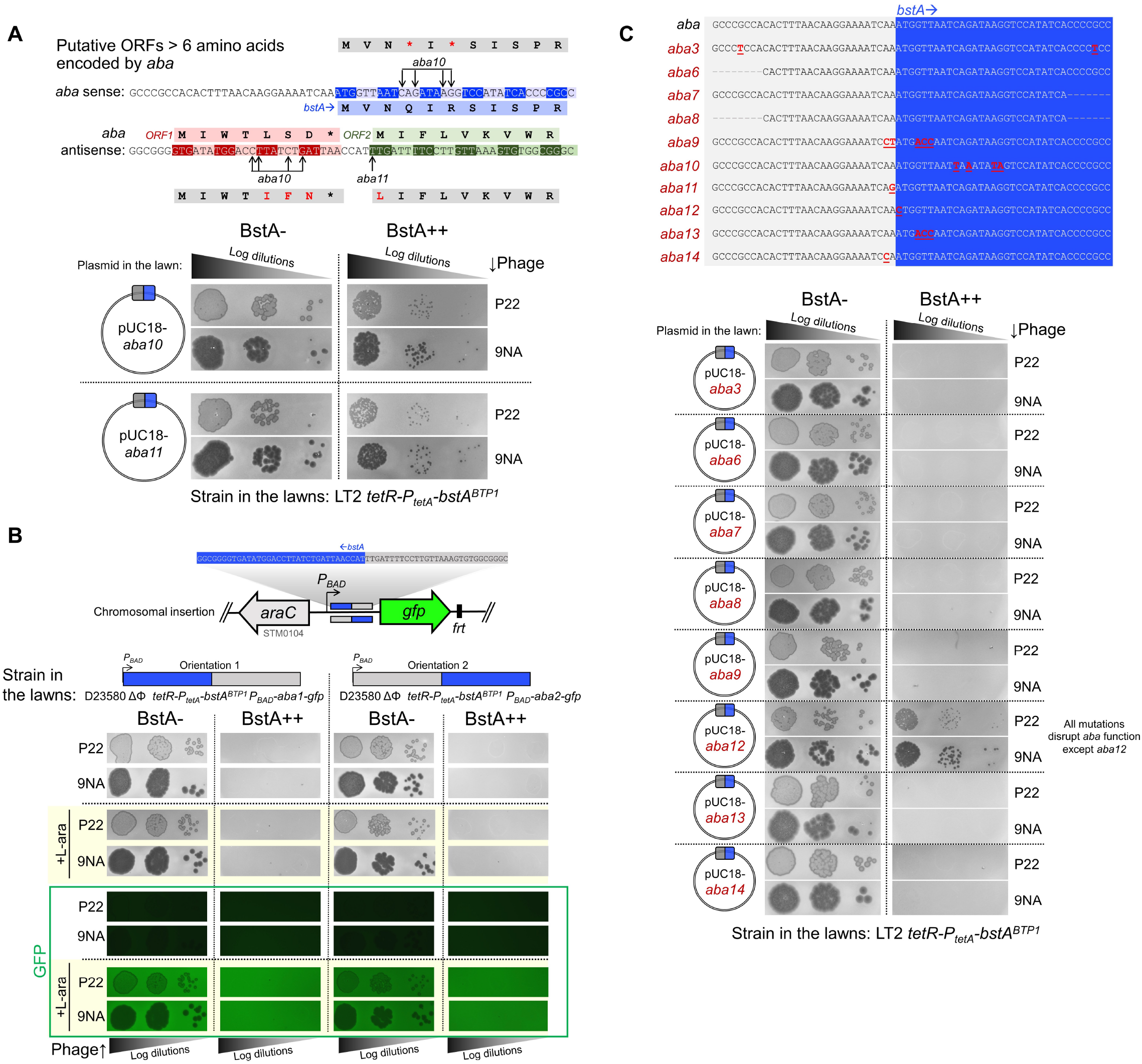
*aba* element functions as a DNA element, and its function is highly sensitive to sequence mutations. (**A**) Analysis of the *aba* sequence: putative open reading frames (ORFs) encoding for peptides longer that 6 amino acids are shown. All the ATG, TTG or GTG triplets were considered as putative start codons. The mutations of the functional *aba10* and *aba11* and the resulting mutated residues (in red) in the putative ORFs are indicated. Stop codons are indicated by *. Non-synonymous mutation in the putative ORFs does not disrupt *aba* suppression of BstA, therefore *aba* is unlikely to function *via* a peptide. (**B**) Transcription of *aba* does not counteract the BstA anti-phage activity. The *aba* sequence followed by a *gfp* reporter gene were inserted in both directions in the chromosome of strain D23580 ΔΦ *tetR-P_tetA_-bstA^BTP1^* (strains SNW645 and SNW646), downstream of the native arabinose-inducible promoter *P_BAD_* (bent arrow) controlled by the AraC regulatory protein. Plaque assays were performed with the indicated phages and strains in the presence or absence of L-arabinose (0.2%). Induction of the transcription of *aba* by L-arabinose did not suppress the activity of BstA (P22 and 9NA phage plaquing is inhibited). Induction of *aba* transcription by the *P_BAD_* promoter was confirmed by detection of fluorescence with addition of +L-ara. (**C**) Further mutational disruption was made to the *aba* sequence cloned into pUC18 plasmids, to identify regions of the sequence that are essential for *aba* function. All mutations to pUC18 *aba* were non-functional (i.e. not capable of BstA suppression) except for the *aba12* derivative. Plaques assays were performed with phage P22 and 9NA applied on lawns of mock-induced (BstA-) or AHT-induced (BstA++) LT2 *tetR-P_tetA_-bstA^BTP1^* (JH4400), transformed with the indicated plasmid.

**Supplementary Figure 9.**
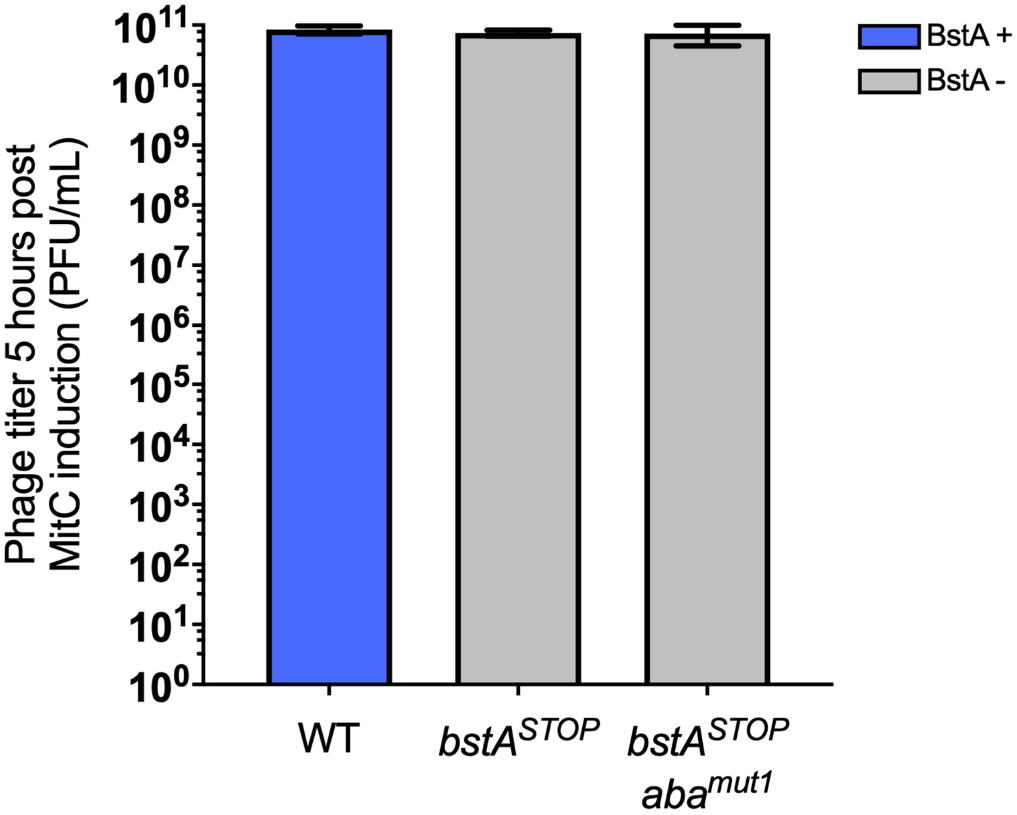
The *aba ^mut1^* mutation has no effect on prophage induction in the absence of BstA protein expression. Prophage induction was measured in strain D23580 ΔΦ lysogenized with BTP1 WT (SSO-127), BTP1 *bstA^STOP^* (SNW553) and BTP1 *bstA^STOP^ aba^mut1^* (SNW660) after 5 hours post induction with Mitomycin C (MitC). Data are presented as the mean of biological triplicates ± SD. Prophage BTP1 WT has a functional BstA protein

## Notes

https://github.com/baymlab/2020_Owen-BstA

### Competing Interest Statement

The authors have declared no competing interest.

## References

Abedon, S.T. (2012). Bacterial ‘immunity’ against bacteriophages. Bacteriophage 2, 50–54.

Bialek-Davenet, S., Criscuolo, A., Ailloud, F., Passet, V., Jones, L., Delannoy-Vieillard, A.S., Garin, B., Hello, S. Le, Arlet, G., Nicolas-Chanoine, M.H., et al. (2014). Genomic definition of hypervirulent and multidrug-resistant *klebsiella pneumoniae* clonal groups. Emerging Infectious Diseases.

Bingham, R., Ekunwe, S.I., Falk, S., Snyder, L., and Kleanthous, C. (2000). The major head protein of bacteriophage T4 binds specifically to elongation factor Tu. J. Biol. Chem. 275, 23219–23226.

Bondy-Denomy, J., and Davidson, A.R. (2014). When a virus is not a parasite: the beneficial effects of prophages on bacterial fitness. J Microbiol. 52, 235–242.

Brüssow, H., Canchaya, C., and Hardt, W.-D. (2004). Phages and the Evolution of Bacterial Pathogens: from Genomic Rearrangements to Lysogenic Conversion. Microbiol Mol Biol Rev 68, 560–602.

Buchan, D.W.A., and Jones, D.T. (2019). The PSIPRED Protein Analysis Workbench: 20 years on. Nucleic Acids Res 47, W402–W407.

Canals, R., Hammarlöf, D.L., Kröger, C., Owen, S.V., Fong, W.Y., Lacharme-Lora, L., Zhu, X., Wenner, N., Carden, S.E., Honeycutt, J., et al. (2019). Adding function to the genome of African Salmonella Typhimurium ST313 strain D23580. PLOS Biology 17, e3000059.

Capek, M., Janácek, J., and Kubínová, L. (2006). Methods for compensation of the light attenuation with depth of images captured by a confocal microscope. Microsc. Res. Tech. 69, 624–635.

Cenens, W., Mebrhatu, M.T., Makumi, A., Ceyssens, P.-J., Lavigne, R., Van Houdt, R., Taddei, F., and Aertsen, A. (2013). Expression of a novel P22 ORFan gene reveals the phage carrier state in Salmonella typhimurium. PLoS Genet. 9, e1003269.

Cherepanov, P.P., and Wackernagel, W. (1995). Gene disruption in *Escherichia coli:* TcR and KmR cassettes with the option of Flp-catalyzed excision of the antibiotic-resistance determinant. Gene.

Chopin, M.-C., Chopin, A., and Bidnenko, E. (2005). Phage abortive infection in lactococci: variations on a theme. Current Opinion in Microbiology 8, 473–479.

Cohen, D., Melamed, S., Millman, A., Shulman, G., Oppenheimer-Shaanan, Y., Kacen, A., Doron, S., Amitai, G., and Sorek, R. (2019). Cyclic GMP-AMP signalling protects bacteria against viral infection. Nature 574, 691–695.

Corcoran, C.P., Podkaminski, D., Papenfort, K., Urban, J.H., Hinton, J.C.D., and Vogel, J. (2012). Superfolder GFP reporters validate diverse new mRNA targets of the classic porin regulator, MicF RNA. Molecular Microbiology.

Cumby, N., Davidson, A.R., and Maxwell, K.L. (2012). The moron comes of age. Bacteriophage 2, 225–228.

Datsenko, K.A., and Wanner, B.L. (2000). One-step inactivation of chromosomal genes in Escherichia coli K-12 using PCR products. Proceedings of the National Academy of Sciences of the United States of America.

Dedrick, R.M., Jacobs-Sera, D., Guerrero Bustamante, C.A., Garlena, R.A., Mavrich, T.N., Pope, W.H., Reyes, J.C.C., Russell, D.A., Adair, T., Alvey, R., et al. (2017). Prophage-mediated defense against viral attack and viral counter-defense. Nat Microbiol 2, 16251.

Degnan, P.H., Michalowski, C.B., Babić, A.C., Cordes, M.H.J., and Little, J.W. (2007). Conservation and diversity in the immunity regions of wild phages with the immunity specificity of phage lambda. Mol. Microbiol. 64, 232–244.

Doron, S., Melamed, S., Ofir, G., Leavitt, A., Lopatina, A., Keren, M., Amitai, G., and Sorek, R. (2018). Systematic discovery of anti-phage defense systems in the microbial pan-genome. Science 359.

Doublet, B., Douard, G., Targant, H., Meunier, D., Madec, J.Y., and Cloeckaert, A. (2008). Antibiotic marker modifications of λ Red and FLP helper plasmids, pKD46 and pCP20, for inactivation of chromosomal genes using PCR products in multidrug-resistant strains. Journal of Microbiological Methods.

Durmaz, E., and Klaenhammer, T.R. (2007). Abortive Phage Resistance Mechanism AbiZ Speeds the Lysis Clock To Cause Premature Lysis of Phage-Infected Lactococcus lactis. Journal of Bacteriology 189, 1417–1425.

Eddy, S.R. (1998). Profile hidden Markov models. Bioinformatics 14, 755–763.

Edwards, R.A., Allen Helm, R., and Maloy, S.R. (1999). Increasing DNA transfer efficiency by temporary inactivation of host restriction. BioTechniques.

El-Gebali, S., Mistry, J., Bateman, A., Eddy, S.R., Luciani, A., Potter, S.C., Qureshi, M., Richardson, L.J., Salazar, G.A., Smart, A., et al. (2019). The Pfam protein families database in 2019. Nucleic Acids Res. 47, D427–D432.

Van Den Ent, F., and Löwe, J. (2006). RF cloning: A restriction-free method for inserting target genes into plasmids. Journal of Biochemical and Biophysical Methods.

Figueira, R., Watson, K.G., Holden, D.W., and Helaine, S. (2013). Identification of *Salmonella* pathogenicity island-2 type III secretion system effectors involved in intramacrophage replication of *S. enterica* serovar Typhimurium: Implications for rational vaccine design. MBio.

Fineran, P.C., Blower, T.R., Foulds, I.J., Humphreys, D.P., Lilley, K.S., and Salmond, G.P.C. (2009). The phage abortive infection system, ToxIN, functions as a protein-RNA toxin-antitoxin pair. Proc. Natl. Acad. Sci. U.S.A. 106, 894–899.

Fortier, L.-C., and Sekulovic, O. (2013). Importance of prophages to evolution and virulence of bacterial pathogens. Virulence 4, 354–365.

Gerlach, R.G., Hölzer, S.U., Jäckel, D., and Hensel, M. (2007). Rapid engineering of bacterial reporter gene fusions by using red recombination. Applied and Environmental Microbiology.

Green, R., and Rogers, E.J. (2014). Chemical Transformation of *E. coli*. Methods Enzymol. 329–336.

Hammarlöf, D.L., Kröger, C., Owen, S. V., Canals, R., Lacharme-Lora, L., Wenner, N., Schager, A.E., Wells, T.J., Henderson, I.R., Wigley, P., et al. (2018). Role of a single noncoding nucleotide in the evolution of an epidemic African clade of Salmonella. Proceedings of the National Academy of Sciences of the United States of America.

Hampton, H.G., Watson, B.N.J., and Fineran, P.C. (2020). The arms race between bacteria and their phage foes. Nature 577, 327–336.

Hansen, E.B. (1989). Structure and regulation of the lytic replicon of phage P1. Journal of Molecular Biology 207, 135–149.

Hautefort, I., Proença, M.J., and Hinton, J.C.D. (2003). Single-Copy Green Fluorescent Protein Gene Fusions Allow Accurate Measurement of *Salmonella* Gene Expression In Vitro and during Infection of Mammalian Cells. Applied and Environmental Microbiology.

Heckman, K.L., and Pease, L.R. (2007). Gene splicing and mutagenesis by PCR-driven overlap extension. Nature Protocols.

Heringa, S., Monroe, J., and Herrick, J. (2007). A Simple, Rapid Method for Extracting Large Plasmid DNA from Bacteria. Nature Precedings.

Herrero-Fresno, A., Wallrodt, I., Leekitcharoenphon, P., Olsen, J.E., Aarestrup, F.M., and Hendriksen, R.S. (2014). The Role of the st313-td Gene in Virulence of Salmonella Typhimurium ST313. PLoS One 9.

Herrero-Fresno, A., Espinel, I.C., Spiegelhauer, M.R., Guerra, P.R., Andersen, K.W., and Olsen, J.E. (2018). The Homolog of the Gene bstA of the BTP1 Phage from Salmonella enterica Serovar Typhimurium ST313 Is an Antivirulence Gene in Salmonella enterica Serovar Dublin. Infect. Immun. 86.

Howard-Varona, C., Hargreaves, K.R., Abedon, S.T., and Sullivan, M.B. (2017). Lysogeny in nature: mechanisms, impact and ecology of temperate phages. The ISME Journal 11, 1511–1520.

Ikeda, H., and Tomizawa, J. (1965). Transducing fragments in generalized transduction by phage P1. Journal of Molecular Biology.

Iyer, L.M., Koonin, E.V., and Aravind, L. (2002). Extensive domain shuffling in transcription regulators of DNA viruses and implications for the origin of fungal APSES transcription factors. Genome Biology 3, research0012.1.

Jiang, W., Bikard, D., Cox, D., Zhang, F., and Marraffini, L.A. (2013). RNA-guided editing of bacterial genomes using CRISPR-Cas systems. Nature Biotechnology.

Kandathil, S.M., Greener, J.G., and Jones, D.T. (2019). Prediction of interresidue contacts with DeepMetaPSICOV in CASP13. Proteins: Structure, Function, and Bioinformatics 87, 1092–1099.

Kingsley, R.A., Msefula, C.L., Thomson, N.R., Kariuki, S., Holt, K.E., Gordon, M.A., Harris, D., Clarke, L., Whitehead, S., Sangal, V., et al. (2009). Epidemic multiple drug resistant *Salmonella* Typhimurium causing invasive disease in sub-Saharan Africa have a distinct genotype. Genome Research.

Kintz, E., Davies, M.R., Hammarlöf, D.L., Canals, R., Hinton, J.C.D., and van der Woude, M.W. (2015). A BTP1 prophage gene present in invasive non-typhoidal *Salmonella* determines composition and length of the O-antigen of the lipopolysaccharide. Molecular Microbiology.

Koskiniemi, S., Pränting, M., Gullberg, E., Näsvall, J., and Andersson, D.I. (2011). Activation of cryptic aminoglycoside resistance in *Salmonella enterica*. Molecular Microbiology.

Kropinski, A.M., Mazzocco, A., Waddell, T.E., Lingohr, E., and Johnson, R.P. (2009). Enumeration of bacteriophages by double agar overlay plaque assay. Methods in Molecular Biology (Clifton, N.J.).

Labrie, S.J., Samson, J.E., and Moineau, S. (2010). Bacteriophage resistance mechanisms. Nature Reviews Microbiology 8, 317–327.

Lauritsen, I., Porse, A., Sommer, M.O.A., and Nørholm, M.H.H. (2017). A versatile one-step CRISPR-Cas9 based approach to plasmid-curing. Microbial Cell Factories.

Lopatina, A., Tal, N., and Sorek, R. (2020). Abortive Infection: Bacterial Suicide as an Antiviral Immune Strategy. Annual Review of Virology 7, null.

Maloy, S.R. (1990). Experimental techniques in bacterial genetics (Boston, MA).

Martínez-García, E., and de Lorenzo, V. (2011). Engineering multiple genomic deletions in Gram-negative bacteria: Analysis of the multi-resistant antibiotic profile of *Pseudomonas putida* KT2440. Environmental Microbiology.

Maxwell, K.L. (2017). The Anti-CRISPR Story: A Battle for Survival. Molecular Cell 68, 8–14.

McClelland, M., Sanderson, K.E., Spieth, J., Clifton, S.W., Latreille, P., Courtney, L., Porwollik, S., Ali, J., Dante, M., Du, F., et al. (2001a). Complete genome sequence of \textlessi\textgreaterSalmonella enterica\textless/i\textgreater serovar Typhimurium LT2. Nature.

McClelland, M., Sanderson, K.E., Spieth, J., Clifton, S.W., Latreille, P., Courtney, L., Porwollik, S., Ali, J., Dante, M., Du, F., et al. (2001b). Complete genome sequence of *Salmonella enterica* serovar Typhimurium LT2. Nature.

Medina, E.M., Walsh, E., and Buchler, N.E. (2019). Evolutionary innovation, fungal cell biology, and the lateral gene transfer of a viral KilA-N domain. Current Opinion in Genetics & Development 58-59, 103–110.

Meeske, A.J., Nakandakari-Higa, S., and Marraffini, L.A. (2019). Cas13-induced cellular dormancy prevents the rise of CRISPR-resistant bacteriophage. Nature 570, 241–245.

Needleman, S.B., and Wunsch, C.D. (1970). A general method applicable to the search for similarities in the amino acid sequence of two proteins. J. Mol. Biol. 48, 443–453.

Owen, S. V., Wenner, N., Canals, R., Makumi, A., Hammarlöf, D.L., Gordon, M.A., Aertsen, A., Feasey, N.A., and Hinton, J.C.D. (2017). Characterization of the prophage repertoire of African *Salmonella* Typhimurium ST313 reveals high levels of spontaneous induction of novel phage BTP1. Frontiers in Microbiology.

Owen, S.V., Canals, R., Wenner, N., Hammarlöf, D.L., Kröger, C., and Hinton, J.C.D. (2020). A window into lysogeny: revealing temperate phage biology with transcriptomics. Microbial Genomics.

Parma, D.H., Snyder, M., Sobolevski, S., Nawroz, M., Brody, E., and Gold, L. (1992). The Rex system of bacteriophage lambda: tolerance and altruistic cell death. Genes Dev. 6, 497–510.

Pecota, D.C., and Wood, T.K. (1996). Exclusion of T4 phage by the hok/sok killer locus from plasmid R1. J Bacteriol 178, 2044–2050.

Pedulla, M.L., Ford, M.E., Karthikeyan, T., Houtz, J.M., Hendrix, R.W., Hatfull, G.F., Poteete, A.R., Gilcrease, E.B., Winn-Stapley, D.A., and Casjens, S.R. (2003). Corrected sequence of the bacteriophage P22 genome. Journal of Bacteriology.

Potter, S.C., Luciani, A., Eddy, S.R., Park, Y., Lopez, R., and Finn, R.D. (2018). HMMER web server: 2018 update. Nucleic Acids Res 46, W200–W204.

Rigden, D.J. (2002). Use of covariance analysis for the prediction of structural domain boundaries from multiple protein sequence alignments. Protein Eng. 15, 65–77.

Riley, M., Abe, T., Arnaud, M.B., Berlyn, M.K.B., Blattner, F.R., Chaudhuri, R.R., Glasner, J.D., Horiuchi, T., Keseler, I.M., Kosuge, T., et al. (2006). *Escherichia coli* K-12: A cooperatively developed annotation snapshot-2005. Nucleic Acids Research.

Rist, M., and Kertesz, M.A. (1998). Construction of improved plasmid vectors for promoter characterization in Pseudomonas aeruginosa and other gram-negative bacteria. FEMS Microbiol. Lett. 169, 179–183.

Rostøl, J.T., and Marraffini, L. (2019). (Ph)ighting Phages: How Bacteria Resist Their Parasites. Cell Host Microbe 25, 184–194.

Rotman, E., Amado, L., and Kuzminov, A. (2010). Unauthorized horizontal spread in the laboratory environment: The tactics of Lula, a temperate lambdoid bacteriophage of *Escherichia coli*. PLoS ONE.

Sambrook, J., and Russell, D.W. (2001). Molecular Cloning: A Laboratory Manual, Third Edition. In Molecular Cloning: A Laboratory a Manual, p.

Samson, J.E., Magadán, A.H., Sabri, M., and Moineau, S. (2013). Revenge of the phages: defeating bacterial defences. Nature Reviews Microbiology 11, 675–687.

Schindelin, J., Arganda-Carreras, I., Frise, E., Kaynig, V., Longair, M., Pietzsch, T., Preibisch, S., Rueden, C., Saalfeld, S., Schmid, B., et al. (2012). Fiji: an open-source platform for biological-image analysis. Nature Methods 9, 676–682.

Schmieger, H. (1972). Phage P22-mutants with increased or decreased transduction abilities. MGG Molecular & General Genetics.

Schulte, M., Sterzenbach, T., Miskiewicz, K., Elpers, L., Hensel, M., and Hansmeier, N. (2019). A versatile remote control system for functional expression of bacterial virulence genes based on the *tetA* promoter. International Journal of Medical Microbiology.

Seemann, T. (2014). Prokka: rapid prokaryotic genome annotation. Bioinformatics 30, 2068–2069.

Shub, D.A. (1994). Bacterial Viruses: Bacterial altruism? Current Biology 4, 555–556.

Simon, R., Priefer, U., and Pühler, A. (1983). A broad host range mobilization system for *in vivo* genetic engineering: Transposon mutagenesis in gram negative bacteria. Bio/Technology.

Snyder, L. (1995). Phage-exclusion enzymes: a bonanza of biochemical and cell biology reagents? Molecular Microbiology 15, 415–420.

Spiegelhauer, M.R., García, V., Guerra, P.R., Olsen, J.E., and Herrero-Fresno, A. (2020). Association of the prophage BTP1 and the prophage-encoded gene, bstA, with antivirulence of Salmonella Typhimurium ST313. Pathog Dis 78.

Thevenaz, P., Ruttimann, U.E., and Unser, M. (1998). A pyramid approach to subpixel registration based on intensity. Trans. Img. Proc. 7, 27–41.

Tiruvadi Krishnan, S., Moolman, M.C., van Laar, T., Meyer, A.S., and Dekker, N.H. (2015). Essential validation methods for *E. coli* strains created by chromosome engineering. Journal of Biological Engineering.

Trasanidou, D., Gerós, A.S., Mohanraju, P., Nieuwenweg, A.C., Nobrega, F.L., and Staals, R.H.J. (2019). Keeping crispr in check: diverse mechanisms of phage-encoded anti-crisprs. FEMS Microbiol Lett 366.

Trinh, J.T., Székely, T., Shao, Q., Balázsi, G., and Zeng, L. (2017). Cell fate decisions emerge as phages cooperate or compete inside their host. Nature Communications 8, 14341.

Tsao, Y.-F., Taylor, V.L., Kala, S., Bondy-Denomy, J., Khan, A.N., Bona, D., Cattoir, V., Lory, S., Davidson, A.R., and Maxwell, K.L. (2018). Phage Morons Play an Important Role in Pseudomonas aeruginosa Phenotypes. Journal of Bacteriology 200, e00189–18.

Valen, D.V., Wu, D., Chen, Y.-J., Tuson, H., Wiggins, P., and Phillips, R. (2012). A Single-Molecule Hershey-Chase Experiment. Current Biology: CB 22, 1339.

Watson, B.N.J., Vercoe, R.B., Salmond, G.P.C., Westra, E.R., Staals, R.H.J., and Fineran, P.C. (2019). Type I-F CRISPR-Cas resistance against virulent phages results in abortive infection and provides population-level immunity. Nature Communications 10, 5526.

Zimmermann, L., Stephens, A., Nam, S.-Z., Rau, D., Kübler, J., Lozajic, M., Gabler, F., Söding, J., Lupas, A.N., and Alva, V. (2018). A Completely Reimplemented MPI Bioinformatics Toolkit with a New HHpred Server at its Core. Journal of Molecular Biology 430, 2237–2243.

Zinder, N.D., and Lederberg, J. (1952). Genetic exchange in *Salmonella*. Journal of Bacteriology 64, 679–699.

